# Mesoscale volumetric light field (MesoLF) imaging of neuroactivity across cortical areas at 18 Hz

**DOI:** 10.1101/2023.03.20.533476

**Authors:** Tobias Nöbauer, Yuanlong Zhang, Hyewon Kim, Alipasha Vaziri

**Affiliations:** Laboratory of Neurotechnology and Biophysics, The Rockefeller University, New York, NY, USA; Department of Automation, Tsinghua University, Beijing, China; The Kavli Neural Systems Institute, The Rockefeller University, New York, NY, USA

## Abstract

Various implementations of mesoscopes provide optical access for calcium imaging across multi-millimeter fields-of-view (FOV) in the mammalian brain. However, capturing the activity of the neuronal population within such FOVs near-simultaneously and in a volumetric fashion has remained challenging since approaches for imaging scattering brain tissues typically are based on sequential acquisition. Here, we present a modular, mesoscale light field (MesoLF) imaging hardware and software solution that allows recording from thousands of neurons within volumes of ⍰ 4000 × 200 µm, located at up to 400 µm depth in the mouse cortex, at 18 volumes per second. Our optical design and computational approach enable up to hour-long recording of ∼10,000 neurons across multiple cortical areas in mice using workstation-grade computing resources.

---

Information flow across cortical areas is a hallmark of higher-level perception, cognition, and the neuronal network dynamics that underlie complex behaviors. Yet tracing this information flow in a volumetric fashion across mesoscopic fields-of-view (FOV), at a cellular resolution and at a temporal bandwidth sufficient to capture the dynamics of genetically encoded calcium indicators (GECIs)^1–6^, i.e., 10–20 Hz, has remained challenging. In the realm of multi-photon microscopy^7^, several cellular-resolution mesoscopes have been presented that reach FOVs measuring up to ∼5 mm in diameter^8–13^ but typically, fast calcium imaging in these designs is constrained to smaller regions-of-interests. More importantly, since the volumetric imaging rate achievable in serial scanning methods scales as the inverse third power of the side length of the imaged volume, scaling up sequential acquisition approaches to mesoscopic volumes has thus far remained limited and highly involved. Scan-free, mesoscopic widefield one-photon imaging approaches on the other hand, often based on low-NA or photographic objectives^14–16^, have only provided coarse, low-resolution activity information, resolve only superficial neurons^17, 18^, or require sparse expression of GECIs and their targeting to superficial brain regions^19^ to achieve neuron-level discrimination.

In light field microscopy (LFM)^20–28^, a microlens array is used to encode volumetric information about the sample onto a 2D camera sensor. These sensor images are subsequently computationally reconstructed using the system’s point-spread-function (PSF) to obtain 3D sample information. By doing away with the need for scanning excitation, these techniques offer the unique capability to scale up the acquisition volume both laterally and axially without sacrificing frame rate and thus, in principle, can enable fast mesoscopic volumetric imaging. However, due to the limitations imposed by scattering tissues and the computational cost of large-scale deconvolutions, the use of LFM has been restricted to only sub-millimeter FOVs and weakly scattering specimen.

We have recently extended LFM into the scattering mammalian brain^29, 30^ by exploiting the strongly forward-directed nature of light scattering in brain tissue and the capability of LFM to capture both angular and lateral position information contained in the incoming light field. Our Seeded Iterative Demixing (SID) approach^29, 30^ was designed to capture the remaining directional information present in the scattered light-field and, together with the spatio-temporal sparsity of neuronal activity, exploit this information to seed a machine learning algorithm that provides an initial estimate of the locations of the active neurons. SID then iteratively refines both the position estimates and the neuronal activity time series, thereby allowing for neuron localization and extraction of activity signals from depths up to ∼400 µm in the mouse brain.

LFM’s simplicity and scalability combined with SID’s potential to extend this approach into scattering brain tissues in principle makes LFM highly attractive for mesoscale volumetric recording of neuroactivity. However, actual experimental realizations of mesoscopic LFM imaging in the mammalian cortex have thus far been hampered by a lack of solutions for capturing mesoscopic fields-of-views at high optical resolution across multi-millimeter FOVs and appropriate computational tools. On one hand the required computational resources for such tools need to efficiently scale with the imaged volume size – and hence the number of recorded neurons – as well as the number of the recorded frames. On the other hand, these computational tools have to be able to address the unique challenges associated with faithful localization and extraction of neuronal signals at such scale.

High-resolution imaging across multi-millimeter FOVs requires careful correction of optical aberrations, in particular spherical aberration, which scales with the fourth power of the FOV radius and the sufficient correction of which often involves compromises in the correction of other optical aberrations. The computational reconstruction pipeline on the other hand, aside from being able to robustly extract remaining spatial and directional information from the ballistic and scattered photons, has to be able to account for varying tissue morphology and a range of different conditions, such as blood vessels and their pulsation and other sources of non-rigid tissue deformation, while keeping computational cost at bay despite terabyte-scale raw data sizes.

## RESULTS

Here we demonstrate a volumetric, one-photon-based approach that overcomes these challenges. Using a modular, Mesoscale Light Field (MesoLF) imaging hardware and software solution that combines mesoscale optical design and aberration correction with a scalable computational pipeline for neuronal localization and signal extraction we demonstrate volumetric recording from more than 10,500 active neurons across different regions of the mouse cortex within different volumes of ⍰ 4000 × 200 µm positioned at depths up to ∼400 µm. We captured the activity of these neurons at 18 volumes per second and over timespans exceeding one hour per session for which a single workstation equipped with three Graphics Processing Units (GPUs) was sufficient to perform signal extraction and demixing in a matter of hours.

We have designed the MesoLF optical system to be compatible with a widely used commercial mesoscopy platform^31, 32^ which is designed for multiphoton scanning microscopy but lacks well-corrected wide-field imaging capabilities. The MesoLF optical path (Methods, Supplementary Note 1) is based on a custom tube lens consisting of three doublet elements in a configuration akin to the Petzval objective design form^33^. The elements were numerically optimized to correct the output of the mesoscope objective to achieve diffraction-limited imaging of a ⍰ 4-mm-FOV at NA 0.4 and 10× magnification in the 515-535 nm emission range of the GCaMP calcium indicators. Our tube lens design offers a widefield (pre-LFM) optical resolution of ∼600 line-pairs per millimeter across the entire FOV, thus enabling a wide range of high-resolution mesoscopy applications other than LFM, which are often limited by insufficient resolution in large-FOV optics.

To facilitate LFM recording, we placed a microlens array into the image plane of our custom-designed tube lens. An 80-Megapixel CMOS camera captures the resulting LFM raw images at 18 frames per second. All optical components of the MesoLF system, including the 470 nm LED illumination arm, form a module that was integrated into the optical path of our mesoscope via a motorized fold mirror (Methods, Supplementary Note 1, Supplementary Fig. 1).

The MesoLF computational pipeline (Fig. 1a, Methods, Supplementary Notes 2-9, Supplementary Video 1, Supplementary Software 1) is engineered from the ground up to maximize localization accuracy and signal extraction performance at depth in scattering tissue and addresses the challenges associated with scaling the current LFM reconstruction approaches^21, 22^ to mesoscopic volumetric FOVs. Briefly, after tiling the FOV into 6 × 6 patches, correcting for motion, subtracting the global dynamic background, and masking out vasculature pulsation, the MesoLF pipeline generates a temporally filtered activity summary image in which the weakly scattered LFM footprints of active neurons are emphasized relative to the strongly scattered background. A novel phase-space-based LFM deconvolution approach generates a volumetric estimate of the active neuron locations while rejecting fluorescence background from above and below the imaged volume. Subsequent morphological segmentation allows shape-based identification of neuron candidates and their surrounding volumetric neighborhoods (“shells”), and the expected footprints of these neuron- and shell candidates in the LFM raw data are estimated. At the core of the pipeline lies an iterative demixing step in which the spatial and temporal components are alternatingly updated while keeping the respective other fixed. Signals from core and shell components are demixed, and finally, the resulting traces are classified using a convolutional neuronal network. The pipeline is discussed in detail further below, in Supplementary Note 2-9 and Supplementary Fig. 2-10 and illustrated in Supplementary Video 1.

**Figure 1.**
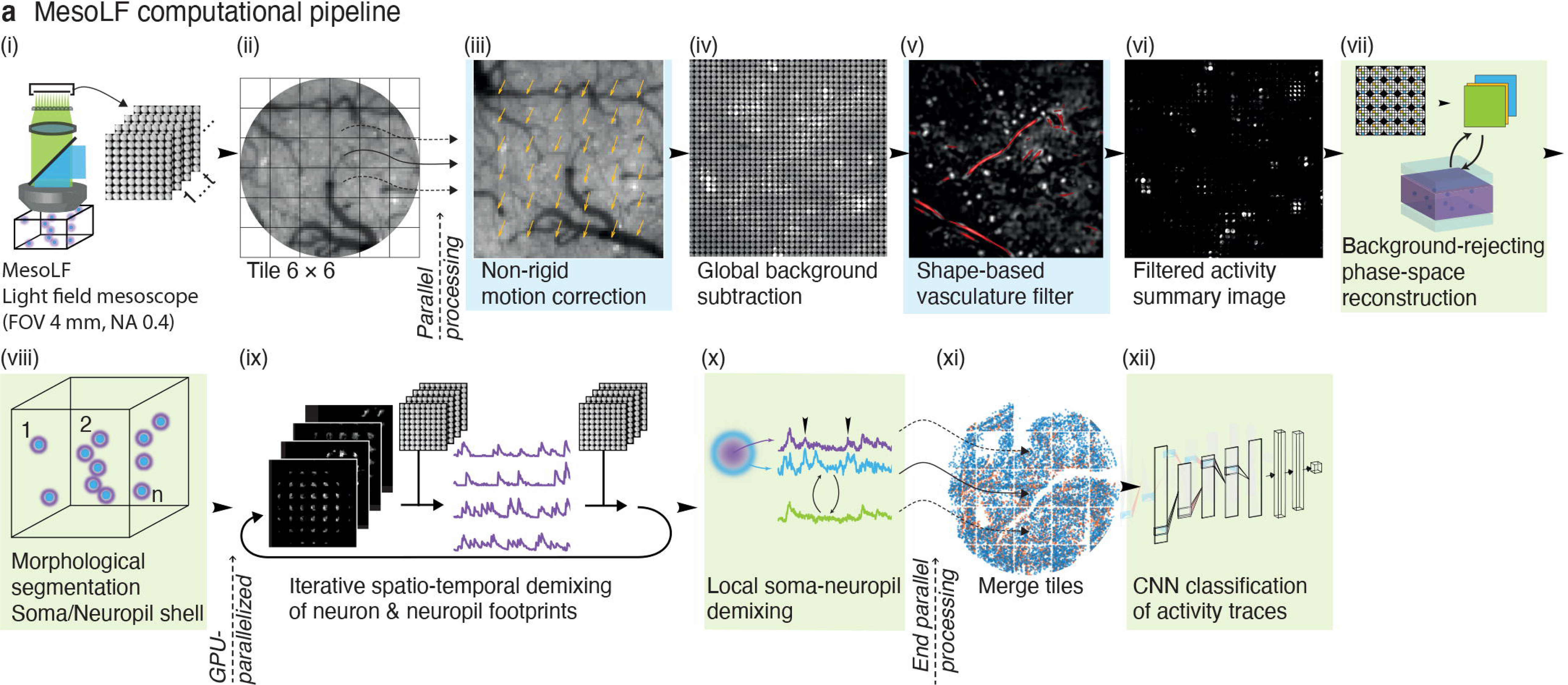
Mesoscopic Light Field (MesoLF) computational pipeline. (a) Schematic overview of key steps of the MesoLF computational pipeline (Main text, Methods, Supplementary Notes 1-9, Supplementary Fig. 1-10, Supplementary Video 1): Fluorescence from the sample is imaged through a custom-designed optical system (FOV ⍰ 4 mm, NA 0.4) and light-field microscope (LFM) detection arm, captured on a CMOS camera (∼ 50M pixels per frame, 6 µm pixel size, 18 fps) and streamed to a flash disk array (i). For offline processing, the frames are tiled into 6 × 6 patches and subsequently processed in parallel on a multi-GPU workstation (ii). Patches undergo non-rigid motion correction (iii) and background subtraction (iv), and blood vessels are detected and masked (v). After temporal filtering of the individual pixel timeseries to remove low- and high-frequency noise, a temporal activity summary image is computed in which temporally active pixels are highlighted (vi). From the summary image, a 3D volume containing the active neurons is reconstructed using a novel artifact-free phase-space reconstruction algorithm that performs background-“peeling”, i.e., the estimation and subtraction of temporally variable background above and below the target volume (vii). A custom morphological segmentation algorithm is applied to segment active neurons in the reconstructed volume (viii). For each candidate neuron and its local surrounding shell, a mask is generated that represents its anticipated spatial footprint in the LFM camera data. In an iterative optimization scheme, these spatial footprints and the corresponding activity time series are updated, thus demixing the neuronal activity signals present in the recording (ix). The resulting neuron- and background-shell signals are further demixed from each other by solving an optimization problem that seeks to reduce crosstalk between neurons and the local background shell components (x). Finally, neuron positions and activity signals from each patch are merged (xi) and classified into high- and low-quality traces by a neuronal network (xii).

We verified the in vivo performance of our high-resolution MesoLF optical module and signal extraction pipeline by performing up to hour-long calcium imaging in the cortex of head-restrained mice. In representative ∼7-minute recordings at 18 Hz (Fig. 2a-c, Supplementary Video 2) of a mouse expressing a modified version of the cell-body-targeted calcium indicator SomaGCaMP7f^34^ (Methods), within a volumetric FOV of ⍰ 4000 × 200 µm^3^ that was positioned at up to 400 um depth, we detected 10,582 active neurons in the depth range of 0–200 µm, 8,076 active neurons in the depth range of 100–300 µm, and 4,746 active neurons in the range of 200–400 µm. The imaged volume contained all or the majority of the posterior parietal, primary somatosensory, primary visual, anteromedial visual, and retrosplenial cortical area. In the extracted temporal signals, clear correlation between bursts of activity and whisker stimulation onsets are observable (white and black triangles in Fig. 2b and Fig. 2c respectively).

**Figure 2.**
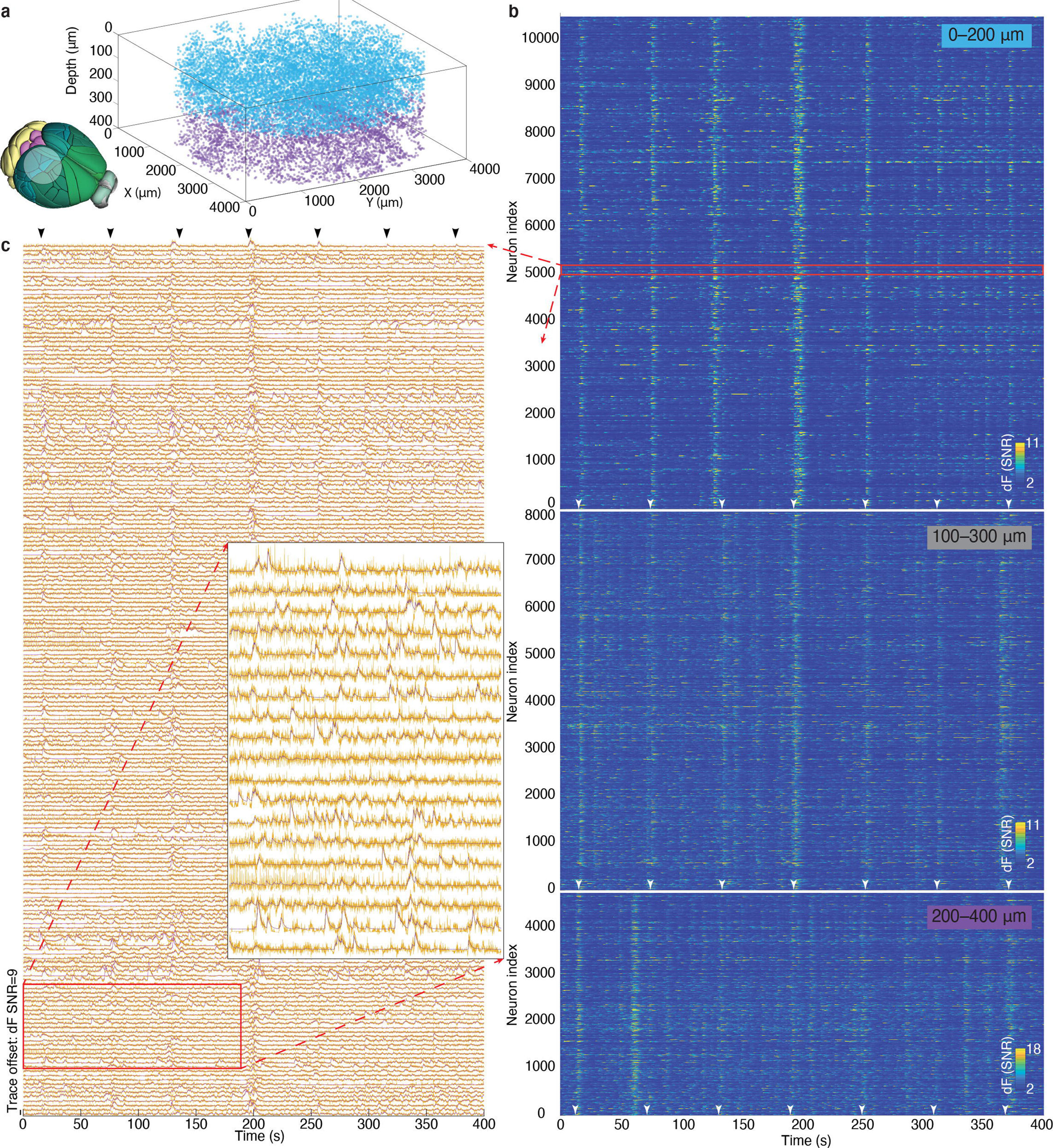
MesoLF calcium imaging in the scattering rodent cortex. **(a)**3D rendering of single neuron positions within an overall volume of ⍰ 4 mm × 400 µm obtained by MesoLF from two subsequent 405-second recordings at 18 volumes per second in mouse cortex. Neuron positions from two sequential recordings of different depth ranges are shown together. Color-coding indicates the record-ing depth range: blue, 0–200 μm; purple, 200–400 μm. Inset: Schematic of imaged field of view superimposed onto a perspective view onto Alan Mouse Brain Reference Atlas. Cortical areas contained in the imaged FOV include the posterior parietal, primary somatosensory, primary visual, anteromedial visual, and retrosplenial cortical area. (b) Heat maps of temporal signals extracted from three 405-second recordings at 18 volumes per second in mouse cortex at three different depth ranges. Neurons are sorted by depth (lower neuron index corresponds to lower depth). Top panel, 10,580 neurons detected in depth range 0–200 µm. Middle panel: 8,076 neurons found in depth range 100–300 µm. Bottom panel: 4,746 neurons found in 200–400 µm depth range. Top and bottom panels correspond to the neuron positions shown in a in blue and violet, respectively. Traces shown are those retained by the CNN trace candidate classifier in “sensitive” mode. Of the traces shown, 1817 (0–200 μm), 1112 (100–300 μm) and 709 (200–400 μm) were classified as high quality and the remainder as intermediate quality. The red rectangle indicates the zoom-in region shown in c. White arrows indicate whisker stimulus onset. Traces represent denoised fluorescence change (dF) normalized to the noise level, defined as the standard deviation of the residuals left after subtracting a low-pass-filtered version of each trace from itself. Color scales are clipped to 15^th^ and 99.9^th^ percentile of all values in each panel for visual clarity. (c) Stacked neuronal activity traces for the region indicated by the red rectangle in b. Traces are normalized by their noise level as in b. Spacing of the traces corresponds to 9 standard deviations of the noise. Yellow lines: un-denoised output of MesoLF pipeline. Violet lines: Fit to un-denoised data with an autoregressive model of calcium indicator response as implemented by the CaImAn package. Black arrows indicate whisker stimulus onset. Inset is zoom into area indicated with red rectangle. Data in (a)–(c) representative of 31 recordings from 6 mice. See also Supplementary Video 2 and Supplementary Fig. 12.

The achievable neuron detection sensitivity, signal extraction quality, and neuron localization accuracy at depth in LFM is ultimately limited by reconstruction artifacts due to scatter-induced aberrations as well as by crosstalk between neurons, neuropil, and background activity above and below the imaged volume. In MesoLF, we have addressed these limitations through the following four key conceptual advances:

First, to reduce reconstruction artifacts that are typical of conventional LFM reconstructions^21, 22^ – in particular those affected by light scattering – without resorting to computationally costly regularization constraints, the input data is transformed into a phase-space representation in which the different angular views of the source volume encoded in an LFM raw image are treated separately and thus can be filtered, weighed, and updated in an optimized sequence^35^ (Fig. 3a). In addition, we introduce a novel “background peeling” algorithm in which fluorescent contributions from above and below the target volume are estimated and subtracted. Such out-of-volume background fluorescence is a key limiting factor of the performance of reconstruction algorithms, which try to explain the observed background signal by allocating it to within- volume features and thus generating reconstruction artifacts. We show that phase-space reconstruction together with background peeling visibly reduces artifacts compared to conventional LFM reconstruction^21, 22^ as well as to the previously published phase space reconstruction approach^35^ (Fig. 3b) and significantly improves the well-known structure similarity index measure (SSIM, see Supplementary Note 5) between reconstruction and ground truth for a depth range of 300–400 µm by 88% while reducing the neuron localization error by 64% (Fig. 3c-e). Furthermore, the neuron identification precision (positive predictive value) is improved by 42% and sensitivity (true positive rate) by 144% (Methods, Supplementary Note 4-5, Supplementary Fig. 5-7).

**Figure 3.**
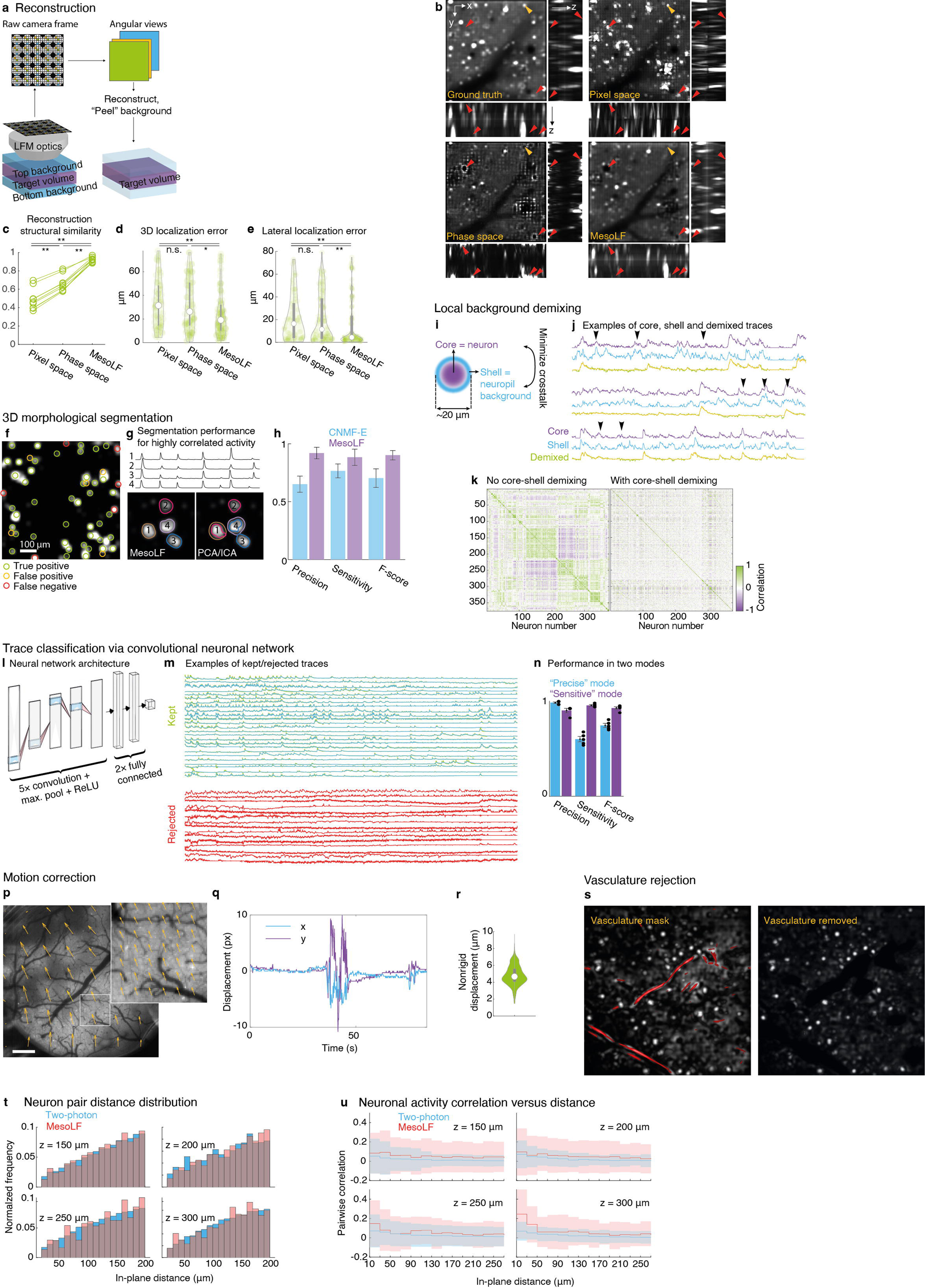
Performance and verification of the individual modules of the MesoLF computational pipeline. (a) Illustration of MesoLF light field phase space reconstruction with background peeling (Supplementary Note 4): Raw camera pixels are re-ordered according to their positions relative to each microlens (colored pixels in raw frame), resulting in a set of angular views (large colored frames) that each represent a perspective from a different angle onto the sample. Reconstruction of the target volume is achieved by iteratively updating the estimate of the volume with weighted and filtered information contained in each of the angular views. To remove artifacts stemming from temporally variable and spatially inhomogeneous background fluorescence from immediately above and below the target volume, the contribution from these top- and bottom background volumes is estimated and subtracted (“peeled”) from the target volume estimate. (b) Comparison of different volumetric light field reconstruction methods and ground truth (see Supplementary Note 4-5). Top left: simulated ground truth volume containing neurons, blood vessels, and neuropil. This volume is numerically convolved with a scattered LFM point spread function to obtain the simulated raw data used as an input for three different reconstruction methods. Top right: Volumetric reconstruction of simulated LFM raw data using Richardson-Lucy deconvolution (Pixel space). Bottom left: Reconstruction using phase space deconvolution without background peeling (Phase space). Bottom right: Reconstruction using phase space deconvolution with background peeling (MesoLF). Red arrows highlight same positions in all panels where artefacts are present in one of the previous methods but absent in MesoLF reconstruction. Yellow arrows indicate position where a ground truth neuron was falsely suppressed in MesoLF. In all panels, the large image is a slice at z = 60 µm in the x-y plane, whereas the smaller images are maximum intensity projections of the reconstructed volume along the x and y axes, respectively. Simulated depth of center of volume: 60 µm. Size of volumes: 600 × 600 × 200 µm^3^, depth range 0–200 µm (c) Structural similarity index between the simulated ground truth volume and the three different classes of reconstructed volumes shown in b (Supplementary Note 5), quantifying quality of reconstruction. n = 9 sets of reconstructions. Paired two-sided Wilcoxon signed rank test for equal median. p = 0.004 (pixel space vs. phase space), 0.004 (pixel space vs. MesoLF), 0.004 (phase space vs. MesoLF). (d) Violin plot of 3D localization error, defined as minimum 3D distance between neurons in simulated ground truth and neurons found in the three different reconstructions shown in b. White circle: median. Thick grey vertical line: Interquartile range. Thin vertical lines: Upper and lower proximal values. Transparent blue disks: data points. Transparent violin-shaped area: Kernel density estimate of data distribution. n = 60, 79, 94 data points, respectively. Two-sided Wilcoxon rank sum test for equal medians, p = 0.567 (pixel space vs. phase space), 0.003 (pixel space vs. MesoLF), 0.019 (phase space vs. MesoLF). n.s., not significant. (e) Violin plot of lateral localization error, defined as minimum lateral distance between neurons in simulated ground truth and neurons found in the three different reconstructions shown in b. Symbols as in d. n = 65, 87, 104 data points, respectively. Two-sided Wilcoxon rank sum test for equal medians, p = 0.766 (pixel space vs. phase space), 0.009 (pixel space vs. MesoLF), 0.029 (phase space vs. MesoLF). n.s., not significant. (f) Segmentation performance in MesoLF (Supplementary Note 6 and Supplementary Fig. 8). Background: Slice from volume reconstruction of temporal summary image from SomaGCaMP7f-labelled mouse cortex, depth 100 µm, simulated data. Colored circles indicate MesoLF segmentation results compared to manual segmentation. (g) Comparison of MesoLF segmentation performance versus PCA/ICA-based segmentation for four simulated neurons with highly correlated temporal activities (activity traces shown above segmented images). Ground truth neurons and corresponding time traces labelled with black digits. Individual segments shown as contour lines with different colors. Note the overlapping and under-segmented output from PCA/ICA. (h) Overall neuron detection scores for the MesoLF morphological segmentation algorithm compared to the CNMF-E package (simulated SomaGCaMP7f-labelled mouse cortex, depth 100 µm). Error bars: Standard deviation. (i) Illustration of core-shell geometry for demixing neuropil activity from soma signals. Signals from segmented regions in f (cores, neurons) and a Gaussian shell region (extending from ∼10 to ∼20 µm diameter) surrounding the cores (background shell, neuropil) are identified and demixed. (j) Sets of representative example traces for core, shell, and demixing result, taken from a recording in mouse cortex at depths 200-400 µm. Arrows indicate crosstalk between shell and core that is removed in the demixed traces. Experimental data from SomaGCaMP7f-labelled mouse cortex, depth 100 µm. (k) Matrices of Pearson correlation coefficients between 400 pairs of neuronal activity traces extracted from a MesoLF recording in mouse cortex, before and after core-shell demixing. The average absolute correlation between signal pairs is reduced by 37% in MesoLF. (l) Illustration of convolutional neural network (CNN) architecture used for classification of candidate neural activity traces (Supplementary Note 9 and Supplementary Fig. 10) (m) Representative examples of 25 kept 10 rejected traces by CNN (Supplementary Note 9 and Supplementary Fig. 10). Traces are experimental data from SomaGCaMP7f-labelled mouse cortex, various depths. (n) Classification performance of two differently trained CNNs, one optimized for a trade-off that prioritizes high precision (“precise mode”, blue bars) and one that prioritizes high sensitivity (“sensitive mode”, violet bars), both while maintaining an overall high F-score. The CNN in “precise” mode achieves precision 0.98 ± 0.01, sensitivity 0.60 ± 0.03, F-score 0.75 ± 0.02; CNN “sensitive” mode achieves precision 0.90 ± 0.02, sensitivity 0.96 ± 0.01, F-score 0.93 ± 0.01. Classification performance was evaluated on withheld data that was not used during training. Error bars: Standard deviation. (Supplementary Note 9 and Supplementary Fig. 10). Black circles: n = 5 data points in each bar. (p) Motion correction in MesoLF. Background is single frame from MesoLF experimental raw data, central sub-aperture image, full FOV (scale bar: 500 µm). Orange arrows indicate direction and magnitude (scaled for clarity, a.u.) of rigid motion correction applied to each of the 6 × 6 tiles into which the raw frame is split at the beginning of the MesoLF pipeline. Inset: Zoom into one of the tiles as indicated with white square. Width of tile: 680 µm. Orange arrows indicate non-rigid motion correction applied within tile. (q) Example of lateral displacement (blue line: x-direction, violet line: y-direction) versus time for one of the tiles in top left panel. (r) Violin plot of non-rigid displacements (i.e., displacements remaining after rigid motion correction). White circle: median. Thick grey vertical line: Interquartile range. Thin vertical lines: Upper and lower proximal values. Transparent blue disks: data points. Transparent violin-shaped area: Kernel density estimate of data distribution. n = 201 data points. (s) Left panel: Example slice from volume reconstruction of MesoLF temporal activity summary image, with vasculature mask overlaid in red. Right panel: Same slice as in left panel, with vasculature removed. Experimental data from SomaGCaMP7f-labelled mouse cortex, depth <50 μm. (t) Comparison of distributions of lateral distances between neuron pairs. Includes all pairs (up to a distance of 200 µm) that can be formed from all neurons in single-plane 2pM recordings (blue bars) at four different depths (150, 200, 250, 300 µm), compared to neurons selected from MesoLF recordings (red bars) at the same depths. Histograms are normalized such that sum of all bin frequencies is one. n = 107,194 neuron pairs in total. (u) Cross-correlation between all pairs of neuronal activity traces versus neuron distance, for all neurons in a 20-µm depth slice centered at four different depths (150, 200, 250, 300 µm) in MesoLF analysis results (red) and 2pM recording at the same depths (blue) (Methods). Solid lines: Median. Shaded areas: Standard deviation. n = 2,230 neurons (MesoLF) and n = 2,282 neurons (2pM) in total.

Second, the implementation of our morphology-based segmentation (Fig. 3f-h, Supplementary Fig. 8, Supplementary Note 6) allows for applying priors on neurons shape and is capable of robustly processing volumes with dense neuron content (Fig. 3f). Compared to the spatio-temporal matrix factorization approaches that have previously been suggested as a way of segmenting active neurons^23, 36^, our purely shape-based approach is not prone to producing segments containing multiple neurons when their temporal activity is highly correlated because it does not rely on temporal independence (Fig. 3g) and overall achieves superior neuron detection performance relative to a comparable one-photon segmentation algorithm^37^ (Fig. 3h). Since segmentation is performed on a reconstruction of a filtered temporal activity summary image, the blurring effects of scattering are strongly suppressed. The reconstruction and segmentation steps have been optimized based on simulations of a realistic optical tissue model (Supplementary Note 3-6, Supplementary Fig. 4-6, Supplementary Fig. 8).

Third, for each of the detected neuron candidates, a spherical shell surrounding the neuron is generated, and both the neurons and shells are convolved with the LFM PSF to generate a library of initial LFM footprints (Fig. 3i). This library of spatial footprint components and associated temporal activity components is then iteratively refined through alternating updates of the spatial and temporal components. Each of the alternated sub-problems is formulated as a so-called LASSO-constrained optimization^38^ which uses a numerical sparsity regularizer to stabilize solutions. The spherical shells are included in the demixing so that they can accommodate the local background that arises from crosstalk from neighboring neurons. After the main demixing stage, these local background contributions are demixed from the neuron activity temporal components through a greedy search approach (Supplementary Note 7, Supplementary Fig. 9). Thereby we could reduce the average absolute correlation between signal pairs by 37% and effectively reject excessive correlations in the extracted signals (Fig. 3j-k).

Finally, to further reject signals arising from non-neuron sources, such as blood vessel pulsation and residual motion, it is beneficial to classify the candidate traces based on whether their temporal activity patterns are compatible with the known response characteristics of GECIs. Several packages exist that allow fitting time series with models of the GECI response^37, 39–41^, but we found them insufficiently selective to robustly reject artefact signals. We therefore designed and trained a convolutional neuronal network (CNN) on a hand-curated dataset in two different modes, one that emphasizes high sensitivity and one that prioritizes precision, both while maintaining overall high F-score (Fig. 3l-m, Supplementary Note 9, Supplementary Fig. 10). Our CNN achieves a classification performance (F-score) of 93% (Fig. 3n, sensitive mode).

Scaling computational functional imaging at neuronal resolution from sub-millimeter to mesoscopic FOVs in the mammalian brain poses unique challenges related to both the intrinsic properties of brain tissue at multi-millimeter scale and the computational scale of the task. Relative displacements due to non-rigid deformation of the brain that arise from animal motion and skull deformations can be as high as ∼10 µm when imaging the brain over multi-millimeter distances. It is therefore imperative to care-fully correct for these deformations to enable mesoscale neuronal resolution imaging. Furthermore, while large blood vessels can usually be avoided in methods covering smaller FOVs, mesoscopic FOVs will always contain a number of large vessels, which cause, if unmitigated, non-rigid deformation and pulsating shadowing effects that will result in false neuronal signals.

In our MesoLF pipeline, we have addressed these challenges as follows: Performing non-rigid motion correction in LFM has previously been hampered by the computational cost of frame-by-frame reconstruction as would be required to make LFM data compatible with established motion correction algorithms. We overcame this limitation by performing non-rigid motion correction on raw LFM data and by transforming them into the so-called phase space representation in which raw image pixels are reordered to form a set of sub-aperture images, each representing a different angular perspective onto the sample. We then corrected for motion and deformations of the phase space slice corresponding to the oblique perspective and applied the same correction to each of the other phase space slices (Fig. 3p-r, Supplementary Fig. 2, Supplementary Note 2).

To avoid artifacts generated by the periodic pulsation of the vasculature, we implement a four-pronged approach (Supplementary Note 8): First, blood vessels are detected and masked based on their tubular shape^42^ (Fig. 3s). Second, all single-pixel time series are filtered to remove low-frequency oscillations originating from pulsation. Third, remaining spatial features that originate from blood vessel motion are rejected during morphological segmentation based on their shapes. Finally, the afore-mentioned CNN-based time series classifier serves to further reject blood vessel artifacts.

To computationally extract neuronal signals and locations from the ∼20 Gigabit/s raw camera data stream would require prohibitively large computational resources if performed on the basis of the conventional LFM frame-by-frame reconstruction method^21, 22^. Our vastly more efficient SID implementation can perform signal extraction on a smaller, 500 × 500 × 200 µm^3^ FOV within few hours on a multi-GPU workstation^29^. In MesoLF, however, the FOV area and hence dataset sizes are ∼64× larger when imaging at the same frame rate. Thus, to enable practical applications of our method, the computational efficiency was significantly enhanced in our MesoLF pipeline. To this end, we devised an accelerated and parallelized scheme that employs a custom GPU-based implementation of the most performance-critical function, a special convolution-like operation required for propagating a light field from the sample to the LFM camera and vice versa (Supplementary Software 1). In addition, the full FOV is sub-divided into 6 × 6 overlapping tiles that can be processed in parallel on multiple GPUs and subsequently merged to avoid duplicate neurons. When compared to the current release of our SID algorithm^29^ (which already requires three orders of magnitude less computation time than conventional frame-by-frame reconstruction of LFM recordings^22, 29^), MesoLF achieves a 63% reduction in CPU core-hours and a 95% reduction in GPU runtime or, correspondingly, a 2.7-fold and a 20-fold speedup at the same computational resources while performing a range of qualitatively new functionalities and achieving a quantitatively better performance. MesoLF thus elevates neuron-resolved, fast computational imaging capacity to the mesoscopic scale.

We previously quantitatively verified and established the performance of a number of key reconstruction and signal extraction modules in our pipeline^29, 30^ using simultaneous LFM and functional ground truth recording via two-photon microscopy (2pM). Here we expanded on our previously established verification methodology using hybrid 2pM ground truth – MesoLF recordings and verified the performance of our entire MesoLF reconstruction pipeline in three complimentary ways: First, by comparing statistical properties of our results to independently acquired 2pM data (Methods). Second, and most importantly, by directly validating our results using simultaneously acquired, volumetric functional MesoLF and 2pM ground truth data (Methods, Supplementary Note 10). Third, by evaluating the performance of both, the individual modules as well as that of our entire MesoLF pipeline on highly realistic simulated data informed by cortical morphology and physiology (Supplementary Note 5, Supplementary Figure 4, Supplementary Figure 11, Methods).

To compare statistical properties of extracted neuronal signals, after performing MesoLF imaging, we recorded single-plane time series recordings from the same animal at four different depths (150, 200, 250, 300 µm; 6.4 frames per second, 2 × 2 mm FOV) using standard 2pM, followed by neuronal segmentation using the CaImAn package^39^ (Methods). We compared the pairwise distance distributions for neurons detected in each of the two recording modalities and found a high level of agreement at all mentioned depths (Fig. 3t).

Further, to verify the fidelity of the extracted signals, we examined the cross-correlations between all pairs of neuronal signals versus lateral neuron pair distance for the 2pM-recorded data and the signals from MesoLF recordings at the same depths (Fig. 3u). Only at our greatest recording depth of 300 µm did we find that the median cross-correlation value for neuron pairs tends to increase for pair distances of less than ∼30 µm in MesoLF recordings compared to the 2pM data, indicating MesoLF’s ability to achieve accurate spatial discrimination and neuronal signal extraction compatible with cellular resolution recordings.

However, since comparison of sequentially acquired datasets only allows for conclusions on a statistical level, to perform a direct and quantitative validation of performance and accuracy of the MesoLF pipeline in terms of neuron detection perfor- mance, neuron localization error and fidelity of extracted neuronal signals, it is necessary to simultaneously acquire MesoLF (LFM) data and volumetric functional ground truth data using 2pM. The direct generation of such volumetric hybrid 2pM–MesoLF functional datasets is fundamentally hampered by the planar nature of 2pM imaging. To overcome this limitation, we conceived a strategy for combining series of eight simultaneously acquired planar 2pM–MesoLF functional recordings each to form a total of five volumetric 2pM–MesoLF functional datasets covering the entire depth range of our method. To combine these single-plane hybrid 2pM–MesoLF recordings into volumetric functional datasets, we exploited the 3D nature of LFM acquisition and computationally shifted the axial location of the fluorescent source plane in the LFM raw data via a simple transformation known as refocusing (Methods, Supplementary Note 10). In this way we obtain volumetric functional datasets in which all temporal activity signals were simultaneously acquired in 2pM and LFM. Using the 2pM data, we then established what we henceforth considered the “ground truth” volumetric functional dataset by automated signal extraction using the well-established CaImAn signal extraction package^41^, followed by manual annotation.

Comparing this volumetric functional ground truth dataset to the output of our MesoLF pipeline applied to the simultaneously acquired LFM data from the same volume (Fig. 4a), we found that the well-known performance scores sensitivity (true positive rate), precision (positive predictive value) and F-score (harmonic mean of sensitivity and precision) for neuron detection reach values of 0.79 ± 0.12, 0.78 ± 0.13, 0.74 ± 0.10 (mean ± std. dev.), respectively, across our depth range (100–400 μm) (Fig. 4b), comparable to the performance achieved by state-of-the art signal extraction algorithms such as Suite2p^40^ and CaImAn^41^, applied to planar 2pM data (cf. Ref. 43 for a performance comparison). The mean neuron localization error (Fig. 4d) across all depth slices is 2.9 ± 1.2 µm laterally, and 8.0 ± 4.8 µm axially, indicating very good neuron localization performance. In addition, we investigated the temporal matching of the extracted neuronal traces against our volumetric 2pM functional ground truth and found a mean temporal correlation between MesoLF and ground truth traces of 0.75 ± 0.16 (n = 835) across all depths (Fig. 4c).

**Figure 4.**
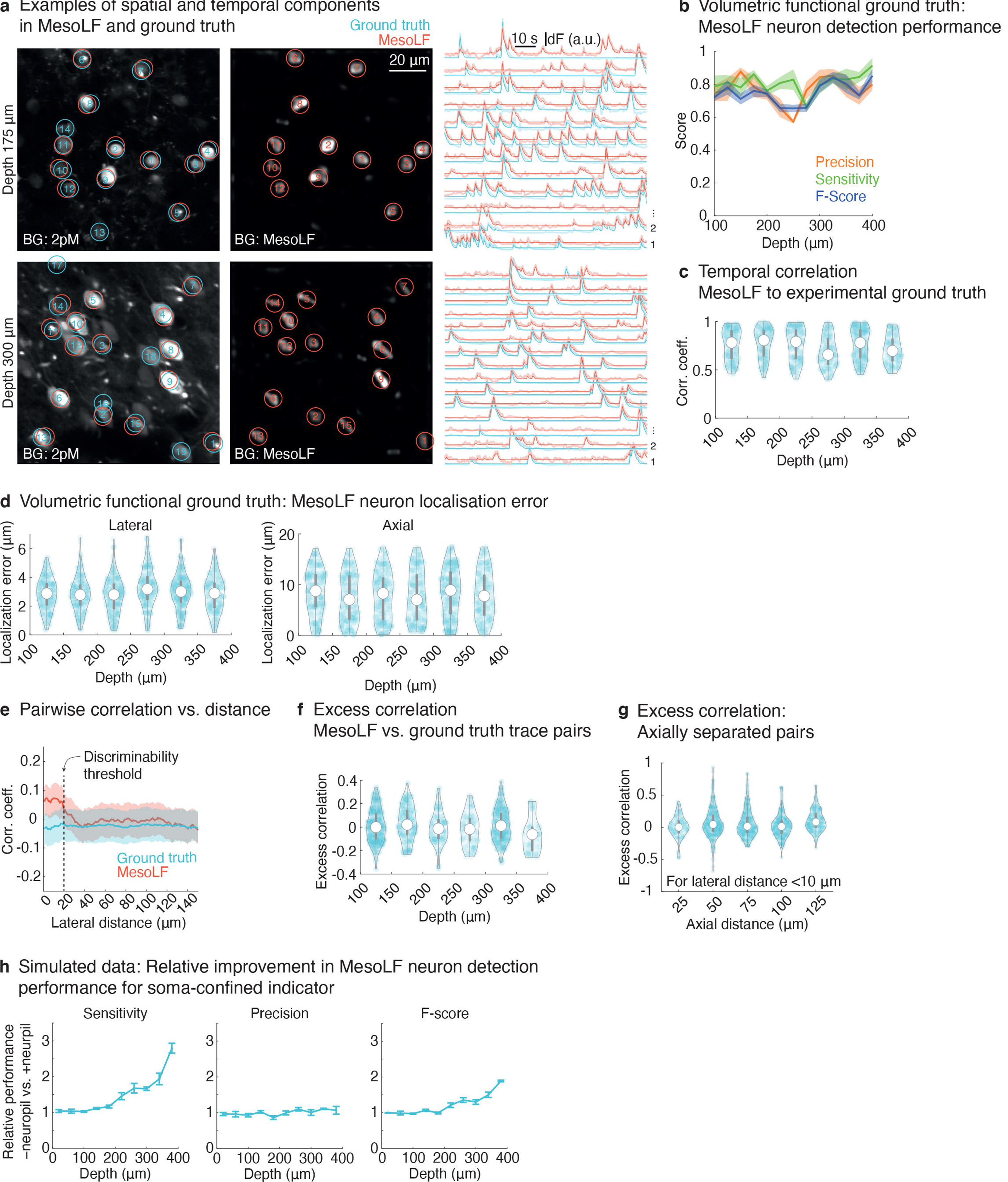
Experimental validation and quantification of performance of full MesoLF pipeline against simultaneously acquired volumetric functional ground truth data. Simultaneously acquired volumetric hybrid 2pM–MesoLF functional ground truth datasets were generated by recording series of planar simultaneous acquisitions in a hybrid 2pM–LFM setup (Methods), followed by refocusing of the axial location of the fluorescent source plane in the LFM raw data and combining eight planar recordings each into volumetric functional datasets (Methods, Supplementary Note 10). Volumetric functional ground truth was established by automatically extracting signals from the 2pM data, followed by human annotation. (a) Left column: Ground truth (blue circles) and MesoLF-extracted neuron positions (red circles) overlaid on a 2pM temporal standard deviation image from a hybrid 2pM– MesoLF recording, for two different example depths (top row: 175 μm; bottom row: 300 μm). Middle column: MesoLF-extracted neuron positions (red circles) overlaid on slice from the reconstructed MesoLF temporal summary frame from the same hybrid 2pM–MesoLF recording as in left column. Right column: Neuronal activity traces corresponding to circles in left and middle column panels, as used for performance quantifications, in experimental functional ground truth (blue traces, corresponding to blue circles in left column panel), recorded by standard 2pM and analyzed using CaI-mAn package followed by human annotation, and simultaneously acquired LFM recordings that were analyzed using the MesoLF computational pipeline (red traces, corresponding to red circles in left column panel), for same two depths as in left column. (b) Neuron detection scores precision, sensitivity and F-score achieved by MesoLF on experimental volumetric functional verification dataset as a function of depth. Shaded areas: mean ± std. dev. n = 80, 77, 69 data points, respectively (c) Distributions of temporal correlations between experimental ground truth activity traces and matched MesoLF traces versus depth. White circle: median. Thick grey vertical line: Interquartile range. Thin vertical lines: Upper and lower proximal values. Transparent blue disks: data points. Transparent violin-shaped area: Kernel density estimate of data distribution. n = 693 data points. (d) Distributions of lateral (left panel) and axial (right panel) neuron localization error between MesoLF-extracted neuron positions and experimental functional ground truth. Violin plot symbols as in c. n = 832,676 data points, respectively. (e) Mean pairwise correlation between all pairs of traces in volumetric experimental functional ground truth (blue line) and mean pairwise correlation between corresponding pairs of MesoLF-extracted traces (red line) as a function of lateral distance between the neurons in the pairs. Only for lateral separations smaller than ∼20 μm, a significant increase in excess correlation, i.e., in the difference between MesoLF-extracted correlation and ground truth correlation, is observable. Shaded areas: Mean ± std. dev. (f) Distribution of excess correlation between pairs of neuronal traces in experimental ground truth and corresponding pairs of MesoLF-extracted traces, as a function of depth. For all depths, the modulus of the median excess correlation is below 0.06, indicating robust crosstalk rejection by the MesoLF pipeline. Violin plot symbols as in c. n = 752 data points. (g) Distributions of excess temporal correlations between pairs of experimental ground truth neuronal activity traces and matched MesoLF traces versus the axial separation of the neuron pair, for neuron pairs with lateral separation < 10 μm. Violin plot symbols as in c. n = 768 data points. (h) For realistically simulated cortical tissue and MesoLF imaging, relative improvement of neuron detection scores achieved by MesoLF when suppressing neuropil labelling, versus depth. The beneficial effect of suppressing neuropil labelling is clearly observable at depths exceeding ∼200 μm. Error bars: Standard error of the mean.

To identify and quantify any artifacts introduced by imperfect demixing of neuronal signals and suppression of background as a function of spatial separation of neurons, we compared the pairwise correlations between pairs of neurons found in ground truth (i.e., physiological correlations) as a function of their lateral and axial distances to the pairwise correlations of the corresponding pairs found in MesoLF-extracted activity traces (Fig. 4e-g). Laterally, only for neuron pair distances smaller than ∼20 μm the mean pairwise correlations in the MesoLF-extracted traces (Fig. 4e) increase significantly (exceed the mean value for large pairwise distances by more than one standard deviation). This pair distance marks the limit down to which MesoLF can faithfully discriminate neuronal signals in the presence of scattering. We note that in the presence of scattering and for functional imaging, it is this “discriminability limit” that determines the effective resolvability of active neuronal signals of MesoLF, and not primarily the purely optical resolution of the light field acquisition system.

To investigate the depth dependence of these pairwise correlations, we computed the excess correlation, defined as the difference in pairwise correlation between ground truth neuron pairs and the corresponding MesoLF neuron pairs, and plot their distributions as a function of depth (Fig. 4f). At all depths, the modulus of the median and the standard deviation of the excess correlation values were below 0.06 and 0.15, respectively, indicating robust demixing and discrimination of neuronal signals by the MesoLF pipeline. We further examined the axial neuron discrimination performance achieved by MesoLF by plotting distributions of excess pairwise correlation as a function of axial distance for neuron pairs with a lateral distance of less than 10 μm, i.e., pairs that are located on top of each other axially (Fig. 4g). This relative location of neurons represents a particularly challenging configuration due to the large spatial overlap in illuminated sensor area that results from two such neurons in the presence of scattering. The standard deviation of the excess correlation was below 0.22 across the axial distances examined, indicating that MesoLF can robustly demix neuronal activity signals even in for this most challenging relative position of two neurons.

The experimental functional ground truth datasets underlying these verification analyses were recorded from animals expressing GCaMP6f throughout the cytosol, i.e., without localization to the soma or the nucleus. This results in significantly less favorable conditions than those that apply to the data shown in our main results (Fig. 2) and as such the functional ground truth results represent a lower bound for the performance of our MesoLF algorithm.

To investigate the contribution of labeled neuropil to our above experimental results and the extent to which MesoLF would benefit from a localized expression of GCaMP, we evaluated MesoLF performance on datasets generated via highly realistic simulations informed by cortical morphology and physiology of neuronal tissue, assuming a neuron density equivalent to the densest regions in our experimental specimen (Supplementary Note 9). These simulations allowed us to selectively disable GECI labelling of neuropil and thus directly compare the effect of using soma-targeted GCaMP to non-localized indicators. The beneficial effect of using soma-targeted GCaMP in combination with MesoLF is apparent from the strong, up to 300% relative enhancement of sensitivity and 200% enhancement of F-score at depth 400 μm (Fig. 4h) in simulations in which labeling of neuropil was absent. Applying MesoLF to simulated datasets with labelled neuropil, we found neuron localization error, temporal activity correla- tions and effects on pairwise correlation structure to be comparable to our experimental results (Supplementary Figure 11), indicating that our simulations and experi- mental ground truth verifications are valid complementary approaches for examining MesoLF performance.

## DISCUSSION

In summary, our MesoLF solution accomplishes mesoscopic high-speed functional imaging of up to 10,500 neurons within volumes of ⍰ 4000 × 200 µm located at up to ∼400 µm depth at 18 volumes per second in the mouse cortex, with only workstationgrade computational resources. This performance is enabled by key advancements in our custom optical design and computational reconstruction and neuronal signal extraction pipeline: Our novel background-rejecting phase-space reconstruction algorithm that is optimized for robustness in the presence of scattering and thus improves reconstruction quality by 88% and reduces the 3D neuron localization error by 64% in tissue simulations. Second, our novel morphological segmentation approach that ef- fectively rejects blood vessel-induced artifacts, does not rely on temporal independence of signals from neighboring neurons and outperforms a comparable one-photon segmentation algorithm^37^ by 22% and 44% in the F-score metric at depths 100 µm and 300 µm, respectively (Fig. 3h; data for depth 300 µm not shown). Third, our core-shell local background demixing solution, which reduces neuropil-neuron crosstalk by 37%. Efficient parallel processing and a custom GPU-accelerated implementation of key processing steps decreases GPU computation time by 95% and CPU core-hours by 63% compared to our previous LFM-based signal extraction solution while boosting the scope, functionality, and performance of signal extraction qualitatively and quantitatively.

By entirely avoiding the inverse cubic scaling relation between volumetric frame rate and side length of the imaging volume that is inherent to serial scanning approaches, the MesoLF concept is uniquely positioned to fully capture the higher temporal band-width (∼500 Hz) offered by genetically encoded voltage indicators^44, 45^, the majority of which are currently optimized for one-photon imaging only, across large volumes in scattering tissue. The achievable frame rate in MesoLF is limited by the number of photons that can be detected per frame while keeping the excitation power and resulting bleaching rate sufficiently low. MesoLF performance will therefore greatly benefit from cameras with improved quantum efficiency, reduced read noise, and faster readout speeds. Here we have shown the performance of MesoLF using GCaMP at up to ∼400 µm depth in the scattering mouse brain, limited by loss of directional information of the scattered photons. The obtainable depths can thus be expected to be further increased in the future by using more efficient and red-shifted indicators.

While several aspects of the MesoLF pipeline are specifically designed to tackle issues arising from large-FOV imaging, the general performance improvements afforded by our implementation will also benefit smaller-scale LFMs, such as our head-mounted MiniLFM device^30^. The MesoLF optical and optomechanical design will be available under an open-source license and the custom tube lens will be commercially obtainable. To facilitate effortless dissemination of our computational pipeline, we will provide a readily deployable container that can be run on common cloud infrastructures, thus lowering the entrance barrier to performing long-duration and high-throughput recording of volumetric calcium activity at mesoscopic FOVs.

## ACKNOWLEDGMENTS

We thank Peer Strogies and James Petrillo at the Rockefeller University’s Precision Instrumentation Technology core facility for mechanical engineering and fabrication of the MesoLF optomechanical components, David Hillebrand for an aliquot of the AAV9-TRE3-2xsomaGCaMP7f virus, Juro Gottweis for an initial GPU-implementation approach of a light-field-related operation, Francisca Martínez-Traub for surgeries, and Frank Tejera and Jeffrey Demas for help with the stimulus/behavioral apparatus and two-photon data acquisition. Research reported in this publication was supported by the National Institute of Neurological Disorders and Stroke of the National Institutes of Health under award numbers 5U01NS103488, 1RF1NS110501, 1RF1NS113251, the National Science Foundation under award number NSF-DBI-1707408 and the Kavli Foundation. T.N. was supported by a Kavli Neural Systems Institute postdoctoral fellowship.

## AUTHOR CONTRIBUTIONS

T.N. designed and implemented the MesoLF optical path under the guidance of A.V., performed experiments, conceptualized and contributed to implementation of the MesoLF computational pipeline, analyzed data, and wrote the manuscript. Y.Z. contributed to conceptualization and implementation of the MesoLF computational pipeline, performed simulations, analyzed data, and wrote the manuscript. H.K. performed cranial window surgeries and viral injections and contributed to the manuscript. A.V. conceived and led the project, conceptualized and guided the imaging, signal extraction and data analysis approach, designed in vivo mouse experiments, and wrote the manuscript.

## METHODS

### Experimental model and subject details

All animal procedures met the National Institutes of Health Guide for Care and Use of Laboratory Animals and were approved by the Institutional Animal Care and Use Committee (IACUC) at The Rockefeller University, New York (protocol number 15848H).

Mice were obtained from The Jackson Laboratory (C57BL/6J) and typically group-housed with a 12h/12h inverted light cycle in standard cages, with food and water ab libitum.

### Virus injection and cranial window surgery

Mice were anesthetized with isoflurane (1–1.5% maintenance at a flow rate of 0.7– 0.9 l/min, RWD Life Science anesthesia machine) and placed in a stereotaxic frame (Kopf Instruments). Dexamethasone (0.4 mg/ml) was administered subcutaneously to manage brain swelling. A ∼1 cm incision was made over the midline of the scalp and the underlying periosteum was cleared from the skull. The scalp was sterilized, then removed after administration of local anesthetic bupivacaine (0.125 mg/ml), and the underlying connective tissue was cleared from the skull. A custom-made stainless-steel head bar was fixed behind the occipital bone with cyanoacrylate glue (Loctite) and covered with black dental cement (Ortho-Jet, Lang Dental). Circular craniotomies (5 mm diameter) were performed over the desired imaging site.

A glass pipette was first back-filled with mineral oil and then front-filled with a genetically expressed calcium indicator adeno-associated virus (AAV9-syn-jGCaMP7s-WPRE; cocktail of AAV9-TRE3-2xsomaGCaMP7f & AAV1-Thy1-tTA; AAV1-hSyn1-GCaMP6f). The pipette was then slowly lowered to each injection site and virus was injected (100–125 nl per site, at 10–25 nl/min; titer 2 × 10^12^–2.6 × 10^13^ vgs/ml) into the brain parenchyma at 200 μm depth (single injection; up to 5 × 5 grid of injections centered at PPC: 2.5 mm AP, 1.8 mm ML, 0.2 mm DV or 0.4 mm DV). During multiple injections, the exposed brain was soaked under cold sterile saline.

After virus injection, a circular 5-mm glass coverslip (#1 thickness, Warner Instruments) was lowered into the craniotomy site and sealed in place with tissue adhesive (Vetbond). The exposed skull surrounding the cranial window was covered with a layer of cyanoacrylate glue and then dental cement.

Post-operative care consisted of 3 days of subcutaneous delivery of meloxicam (0.125 mg/kg), antibiotic-containing feed (LabDiet #58T7), and meloxicam-containing (0.125 mg/tablet) food supplements (Bio-Serv #MD275-M). After surgery, animals were returned to their home cages and were given at least one week for recovery and viral gene expression before being subjected to imaging experiments. Mice with damaged dura or unclear windows were euthanized and were not used for imaging experiments.

Table of animals and preparation parameters

**Table of animals and preparation parameters**
| Mouse ID | Strain | Cranial window diameter | Virus | Injection parameters | Depth ranges & number of recordings |
| --- | --- | --- | --- | --- | --- |
| mfi84 | C57/BL6 | 5 mm | AAV9-TRE3-2xsomaGCaMP7f2<br>AAV1-Thy1-tTA | 1:1 titer ratio, $2.6 \times 10^{13}$ vgs/ml<br>10 injections on grid<br>1000 μm apart<br>100 nl/site, 10 nl/min<br>250 μm depth | 0–200 μm: 1 recording |
| mh150 | C57/BL6 | 5 mm | AAV9-TRE3-2xsomaGCaMP7f2<br>AAV1-Thy1S-tTA | 1:1 titer ratio, $2 \times 10^{12}$ vgs/ml<br>23 injections on grid<br>750 μm apart<br>125 nl per site, 25 nl/min<br>200 μm depth | 0–200 μm: 4 rec.<br>100–300 μm: 6 rec.<br>200–400 μm: 2 rec.<br>300–500 μm: 1 rec. |
| mh155 | C57/BL6 | 5 mm | AAV9-TRE3-2xsomaGCaMP7f2<br>AAV1-Thy1S-tTA | 1:1 titer ratio, $2 \times 10^{12}$ vgs/ml<br>25 injections 750 μm apart<br>5 × 5 grid<br>125 nl per site, 25 nl/min<br>200 μm depth | 100–300 μm: 1 rec.<br>200–400 μm: 2 rec. |
| mh159 | C57/BL6 | 5 mm | AAV9-syn-jGCaMP7s-WPRE | $2 \times 10^{12}$ vgs/ml<br>25 injections 750 μm apart<br>5 × 5 grid<br>125 nl per site, 25 nl/min<br>200 μm depth | 0–200 μm: 3 rec.<br>100–300 μm: 4 rec.<br>200–400 μm: 2 rec. |
| mh160 | C57/BL6 | 5 mm | AAV9-syn-jGCaMP7s-WPRE | $2 \times 10^{12}$ vgs/ml<br>25 injections 750 μm apart<br>5 × 5 grid<br>125 nl per site, 25 nl/min<br>200 μm depth | 0–200 μm: 1 rec |
| mh161 | C57/BL6 | 5 mm | AAV9-syn-jGCaMP7s-WPRE | $2 \times 10^{12}$ vgs/ml<br>25 injections 750 μm apart<br>5 × 5 grid<br>125 nl per site, 25 nl/min<br>200 μm depth | 0–100 μm: 1 rec.<br>100–300 μm: 2 rec.<br>200–400 μm: 1 rec. |
| **Hybrid 2pM-MesoLF functional ground truth verification recordings:** |  |  |  |  |  |
| mo4 | C57/BL6 | 5 mm | AAV1-hSyn1-GCaMP6f | $10^{12}$ vgs/ml<br>2 × 2 injections, 400 μm apart<br>25 nl per site, 10 nl/min<br>400 μm depth | 100–400 μm: 1 rec |
| x01 | C57/BL6 | 5 mm | AAV1-hSyn1-GCaMP6f | $10^{12}$ vgs/ml<br>2 × 2 injections, 400 μm apart<br>25 nl per site, 10 nl/min<br>400 μm depth | 100–300 μm: 1 rec |

### In vivo Ca^2+^ imaging with MesoLF optical system

For MesoLF imaging, animals were head-fixed on a home-built treadmill underneath the HHMI Janelia/Thorlabs 2p-RAM mesoscope objective. The headbar clamp pair was mounted on a two-axis goniometer stage for precision tip/tilt adjustment. Using this goniometer and the 2p-RAM motorized gantry axes (x, y, z, tilt), the cranial window was adjusted to be orthogonal to the optical axis of the 2p-RAM objective. This was achieved using a home-built alignment tool that can be placed into the objective mount and provides a laser reflex from a reference glass plate that is used as the target for aligning the laser reflex from the cranial window.

The MesoLF optical system used for Ca^2+^ imaging is described in detail in Supplementary Note 1 and Supplementary Fig. 1. Briefly, for MesoLF imaging, a motorized fold mirror was moved into the 2p-RAM emission arm to direct fluorescence towards our custom-built MesoLF path and also reflect incoming one-photon excitation light from the MesoLF path towards the 2p-RAM objective.

The MesoLF excitation path consists of a mounted blue LED (Thorlabs M470L3, 470 nm center wavelength, 650 mW), adjustable asphere collimator (Thorlabs SM2F32-A), an iris aperture for adjusting illumination NA, excitation filter (Chroma ET470/40x, ⍰ 2"), engineered diffuser for creating a flat-top intensity profile (RPC Photonics EDC-10-15027-A 2S, 2" square), relay lens (Edmund 45-418, f=300, ⍰ 3") and three fold mirrors. This arrangement provides telecentric, homogeneous illumination in the focal plane in the sample. Illumination power was ∼15 mW post-objective, which corresponds to ∼1.2 mW/mm^2^, a value comparable to our previous LFM imaging methods and typical wide-field imaging protocols.

The MesoLF emission path consists of an emission filter (Semrock Brightline FF01-525/39, ⍰ 2"), microlens array (RPC Photonics MLA-S100-f12, square grid, pitch 100 µm, f = 1.2 mm, F-number 12.5, diced to 42 × 42 mm) and camera (Teledyne DALSA Falcon 4-CLHS 86M, 86 Megapixels, 6 µm pixel pitch, 12 bit, global shutter, 16 fps full frame rate). The excitation and emission paths are combined using a dichroic beamsplitter (Semrock FF505-SDi01 short-pass dichroic, 80 × 50 mm).

Both excitation and emission pass through a custom-designed tube lens (Supplementary Note 1, Supplementary Fig. 1) that corrects aberrations left uncorrected by the 2p-RAM objective in the visible range to achieve diffraction-limited resolution at NA 0.4 in the GCaMP-compatible emission window at 515-535 nm.

For two-photon imaging, the motorized fold mirror mentioned above was moved out of the 2p-RAM detection path so that the system was operating as designed in two-photon imaging mode. Two-photon data was analyzed using the CaImAn signal extraction package^41^.

### Apparatus for stimulus delivery and behavioral tracking

Visual and somatosensory stimuli were controlled via a pre-programmed pulse table generated by National Instruments DAQ cards in the experiment control PC. For whisker stimulation, an Arduino microcontroller with a motor shield and servo motor were employed to move a brush forward and backward over the animals’ whiskers at time intervals indicated by the stimulation protocol. The brush size and its proximity were chosen to stimulate all whiskers simultaneously (as opposed to stimulation of specific whiskers), and stimulation was applied contralaterally to the hemisphere being recorded by the microscope.

All rodents were head-fixed on a home-built treadmill with a rotation encoder affixed to the rear axle (Broadcom, HEDS-5540-A02) to measure the relative position of the treadmill during the recordings. Treadmill position, the microcontroller clock value, and the onset of a whisker stimulus were streamed to the control computer via a serial port connection and logged with a separate data logging script. The data logging script also read out frames from a camera (Logitech 860-000451) in order to capture additional animal behavior during recordings. Motion energy (Supplementary Fig. 12) for manually defined regions of interest (e.g., front paws, nose tip) were computed from the behavior videos using the Facemap Python package^46^ as the magnitude of the difference between each frame and a blockwise mean frame.

### Data management and signal extraction using MesoLF computational pipeline

Data was acquired from the camera onto a control workstation (Intel Xeon W-2155 CPU 3.30 GHz, 10 cores, 256 GB RAM, Windows 10) configured with two software-defined RAID-0 arrays of two PCIe flash disks each (2× Samsung 970 EVO 2 TB and 2× Sabrent Rocket 2280 4 TB, respectively) using a custom data acquisition application written in VisualC# .NET. The magnified image covers an area of IZ40 mm on the camera, which corresponds to ∼7000 × 7000 pixels. This subset of pixels can be read out at 18 fps, resulting in a raw data rate of ∼1320 MB/s.

At the end of each imaging session, the raw data was transferred via 10 Gbit/s network links to a network-attached storage server (Synology RS3618xs).

The MesoLF computational pipeline was run on a multi-GPU workstation (Titan Computers) equipped with two Intel Xeon Gold 6136 3.00GHz CPUs with 12 cores each, 260 GB RAM, three nVidia TITAN V GPUs with 12 GB RAM each, a 1 TB NVMe SSD hard disk, two 1 TB SATA SSD hard disks in a RAID-0 configuration, a 10 Gbit/s network card. Xubuntu 20.04 was used as the operating system and all data analysis was performed in MATLAB R2020a (The Mathworks).

Running the MesoLF analysis shown in Fig. 2b (7-minute recording, 18 fps) took a total of 316 CPU core-hours and 4.1 GPU-hours, as tracked using the pidstat command. This includes loading the raw data from the network-attached storage server, which accounts for approx. 20% of the total run time and can be accelerated by holding data on local SSD disks. The full analysis run was completed within 23 hours and 26 minutes.

### Hybrid 2pM–MesoLF functional ground truth recordings

Direct verification of MesoLF compared to a ground truth is inherently limited by the comparably low voxel rates available in established methods such as 2pM, and due to the planar nature of 2pM. The hybrid 2pM–MesoLF functional ground truth recordings used for performance validation of the MesoLF computational pipeline were recorded on a custom hybrid 2pM–LFM microscope. The instrument is based on Scientifica Slicescope 2pM platform with a custom LFM detection arm. Two-photon excitation pulses (920 nm, 140 fs pulse duration, 80 MHz pulse rate, Coherent Chameleon) were focused into mouse cortex and scanned in planes parallel to the cranial window at a series of depths, via the Slicescope’s galvo-galvo scan path and a Nikon 16×/0.8NA objective mounted on a motorized stage that allowed for axial translation.

Fluorescence from the sample was split at a 10:90 ratio between the Slicescope’s non-descanned PMT arm (emission filter: 525/50 nm, GaAsP PMT, Hamamatsu) and a custom-built LFM arm using a 10% beam sampler (Omega) inserted behind the objective. For LFM detection, fluorescence passed through the short-pass dichroic that couples the laser into the beam path, as well as a GFP emission filter. The image formed by a standard Olympus tube lens was then relayed via two 2-inch achromat lenses (f = 200 mm, Thorlabs) onto a microlens array (MLA, Okotech, custom model, size 1" square, f-number 10, 114 µm microlens pitch, quadratic grid, no gaps). The f-number of the MLA was matched to the output f-number of the microscope. The back focal plane of the MLA was relayed by a photographic macro objective (Nikon 105 mm/2.8) at unity magnification onto the sensor of an Andor Zyla 5.5 sCMOS scientific camera (2560 × 2160 px, 16 bit). To introduce an offset between the 2pM focal plane and the LFM native focal plane, the MLA and camera were translated backwards by a distance corresponding to 40 μm in sample space. The 2pM frame clock was used to trigger camera exposures. A FOV of 200 × 200 µm was scanned at a frame rate of 5 Hz. 2- minute movies were recorded both in the PMT and the LFM camera channel at 13 depths in steps of 25 μm, ranging from 100 to 400 μm in mouse cortex.

In the LFM raw data recorded in this way, fluorescence appears to be emanating from an axial plane offset 40 μm from the axial center of the LFM volumetric field of view (due to the aforementioned displacement of LFM camera and MLA backwards from the rear focal plane of the microscope).

To combine these single-plane hybrid 2pM–MesoLF recordings into volumetric functional datasets, we exploited the 3D nature of LFM acquisition and computationally shifted the axial location of the fluorescent source plane in the LFM raw data via a simple transformation known as refocusing^47^ (Supplementary Note 10). With the a priori knowledge that all light in the LFM raw data came from a single axial plane of known depth with respect to the native focal plane, this transformation is unambiguous, relies only on elementary properties of LFM imaging and treats ballistic and scattered light in an unbiased manner. This approach allowed us to refocus and add 8 single-plane recordings such that they result in a single dataset that contains fluorescence emanating from throughout the entire LFM volumetric FOV of 200 μm axially. We built such volumetric LFM movies for two depth ranges, 100–300 and 200-400 μm, for each recorded session.

The synthetic volumetric LFM functional datasets were then processed using the MesoLF pipeline. The 2pM data was analyzed plane by plane using the CaImAn package, followed by human annotation of the CaImAn results (removing false positives, adding false negatives). This resulted in the set of neuronal positions and activity time traces that was subsequently considered the ground truth.

The MesoLF-extracted neuron locations and time traces were then classified as true positives if the centroid of their spatial filter was within 30 μm to the centroid of a ground truth neuron and had a temporal correlation with the ground truth activity trace of > 0.4. The performance scores sensitivity, precision and F-scores were calculated from the resulting true/false positive rates found in the MesoLF data. The set of all matched MesoLF- and ground truth neurons was then further analyzed to obtain the distributions for localization errors, temporal correlation to ground truth, and excess correlations between pairs of traces presented in Fig. 4. Equivalent analyses were performed on the data simulated using the NAOMi package with an active neuron density of 14,000 per mm^3^ (Supplementary Note 5) to yield the performance quantifications shown in Fig. 4 and Supplementary Fig. 11.

## DATA AVAILABILITY

The data that support the findings of this study are available from the corresponding author upon reasonable request.

## CODE AVAILABILITY

The custom code that comprises the MesoLF pipeline is available in Supplementary Software. The MesoLF code, which includes including a complete demo script, is available at http://github.com/vazirilab/mesolf. A demo data can be downloaded automatically by the demo script, and is also available at https://doi.org/10.5281/zenodo.7306113.

