## Supplementary Figures for "Mesoscale volumetric light field (MesoLF) imaging of neuroactivity across cortical areas at 18 Hz"

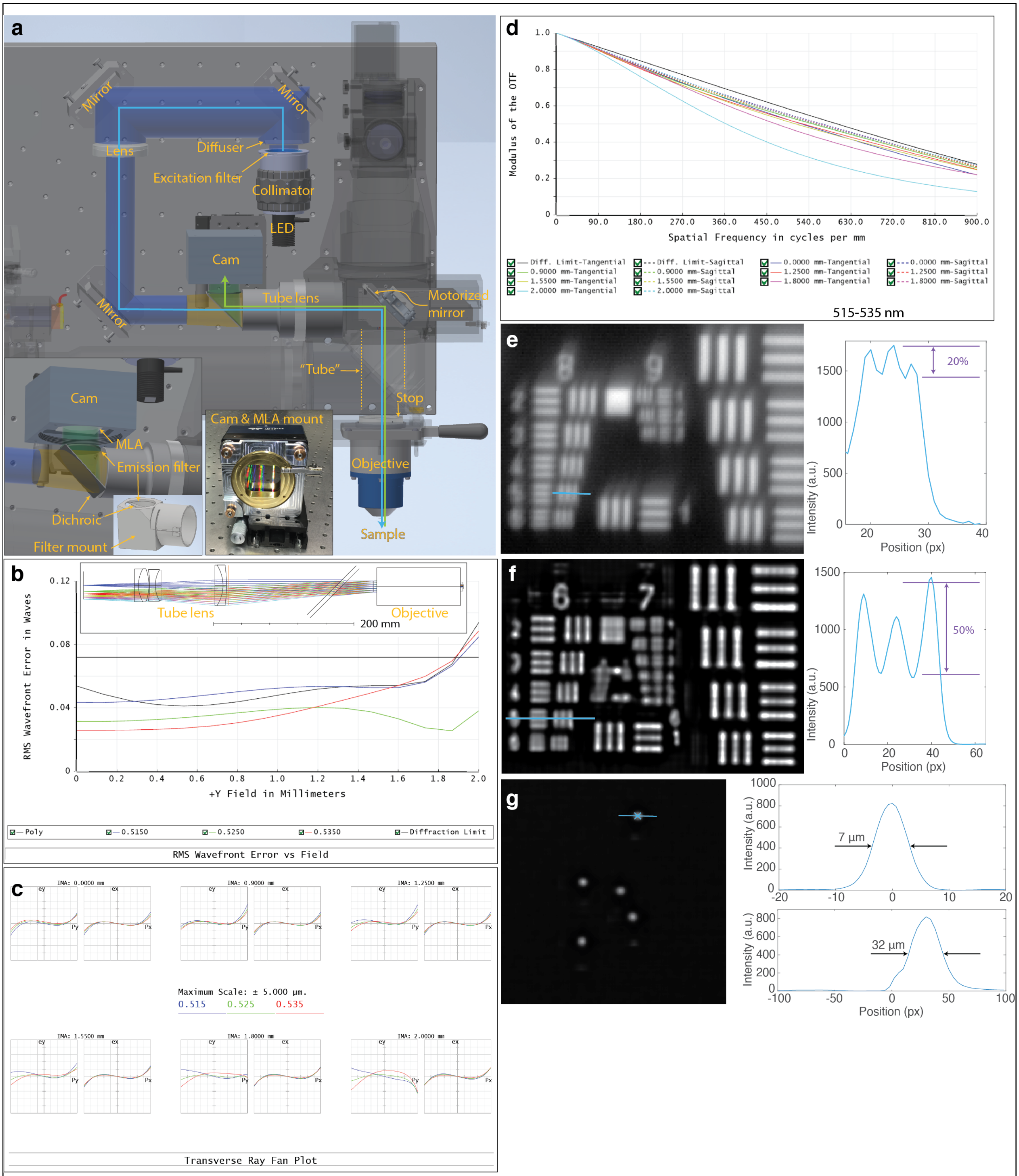


**Supplementary Figure 1**

**MesoLF optical path and optical performance.**

(a) Annotated rendering of main components of the MesoLF optical path. Components of the 2p-RAM system that are not relevant for the MesoLF optical path are shown semi-transparent for clarity. Left inset: Close-up of area around dichroic mirror and camera, not showing dichroic mirror holder, which is instead shown in the sub-inset with filter positions indicated. Right inset: Photo of camera with custom microlens array (MLA) kinematic rotation mount, without MLA installed.

(b) Simulated RMS wavefront error in units of wavelength versus field position, for three different wavelengths (blue solid line: 515 nm, green solid line: 525 nm, red solid line: 535 nm) and as an average over these three wavelengths, weighted to mimic GCaMP excitation spectrum (black solid line). Black horizontal line indicates diffraction limit. Inset: Optical layout of custom tube lens and 2p-RAM objective.

(c) Ray fan plots for 6 different field positions (see labels above panels) and three colors (see legend in center of panel).

(d) Simulated modulation transfer function plotted against spatial frequency in the sample, for six different field positions (line colors, see legend) and the sagittal and tangential planes each (dashed and solid lines, respectively). All resolution lines are averaged over the wavelength range 515-535 nm, weighted to mimic the GCaMP spectrum.

(e) Image of a section of a high-resolution USAF target taken with the MesoLF optical system in wide-field configuration (i.e., no MLA installed, camera in image plane). Right panel: Cross-section along solid line in left panel.

(f) Slice at z = +90 µm from volume reconstruction of MesoLF image of USAF resolution target. Right panel: cross-section along solid line in top panel.

(g) Slice at z = -30 µm from volume reconstruction of MesoLF image of sample containing ⌀6 µm fluorescent beads. Top right: cross-section along solid line in left panel. Bottom right: Axial cross-section through volume along line orthogonal to image plane and through X indicator in left panel.

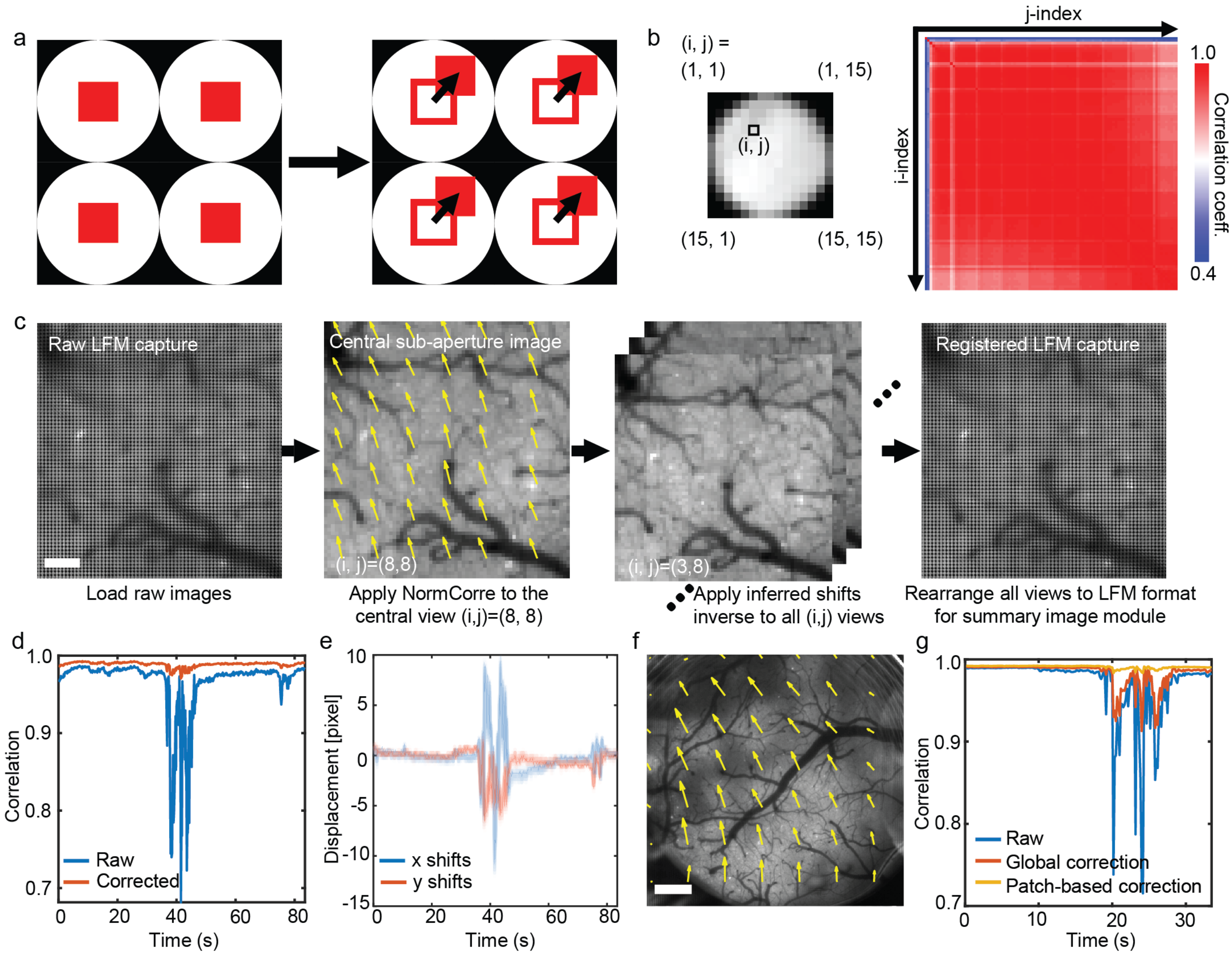


**Supplementary Figure 2**

**Motion correction in MesoLF.**

(a) Illustrated motion patterns in LFM raw data. The lenslet aperture shadows (black areas) will not move, only patterns within those apertures (red squares), which prohibits motion correction using established algorithms without prior rearrangement of the data.

(b) Left panel: LFM image formed behind one microlens, sampled by 15 × 15 pixels. Number pairs in brackets indicate pixel coordinates (i,j). Right panel: Correlation matrix for motion vectors extracted from all 15 × 15 sub-aperture images. Sub-aperture image (i,j) consists of all pixels with coordinates (i,j) relative to the nearest microlens. Consistently high values of correlation across all pairs (i,j) indicate that all sub-aperture images experience similar motion vectors, justifying the use of the same motion correcting transformation across all sub-aperture images.

(c) Illustration of the motion correction pipeline in MesoLF. See Supplementary Notes for narration. Scale bar: 100 µm

(d) Correlation coefficients between successive frames of the central sub-aperture image in an LFM recording of mouse cortical calcium activity, pre- and post-motion correction (blue and red traces, respectively), for a patch size of 660 × 690 µm.

(e) Magnitude of sample motion along lateral axes (x and y, blue and red solid traces, respectively) versus time, for same recording as (d), as extracted using the MesoLF motion correction algorithm. Solid line indicates mean; shaded area indicates standard deviation of motion in different patches across the field of view.

(f) Illustration of motion vectors (arrow size indicates magnitude of motion) for different patches across the 4-mm MesoLF FOV. Scale bar: 500 µm

(g) Correlation coefficients between successive frames of the central sub-aperture movie from the whole MesoLF FOV before motion correction (blue), after global correction (red), and after patch-based correction (yellow).

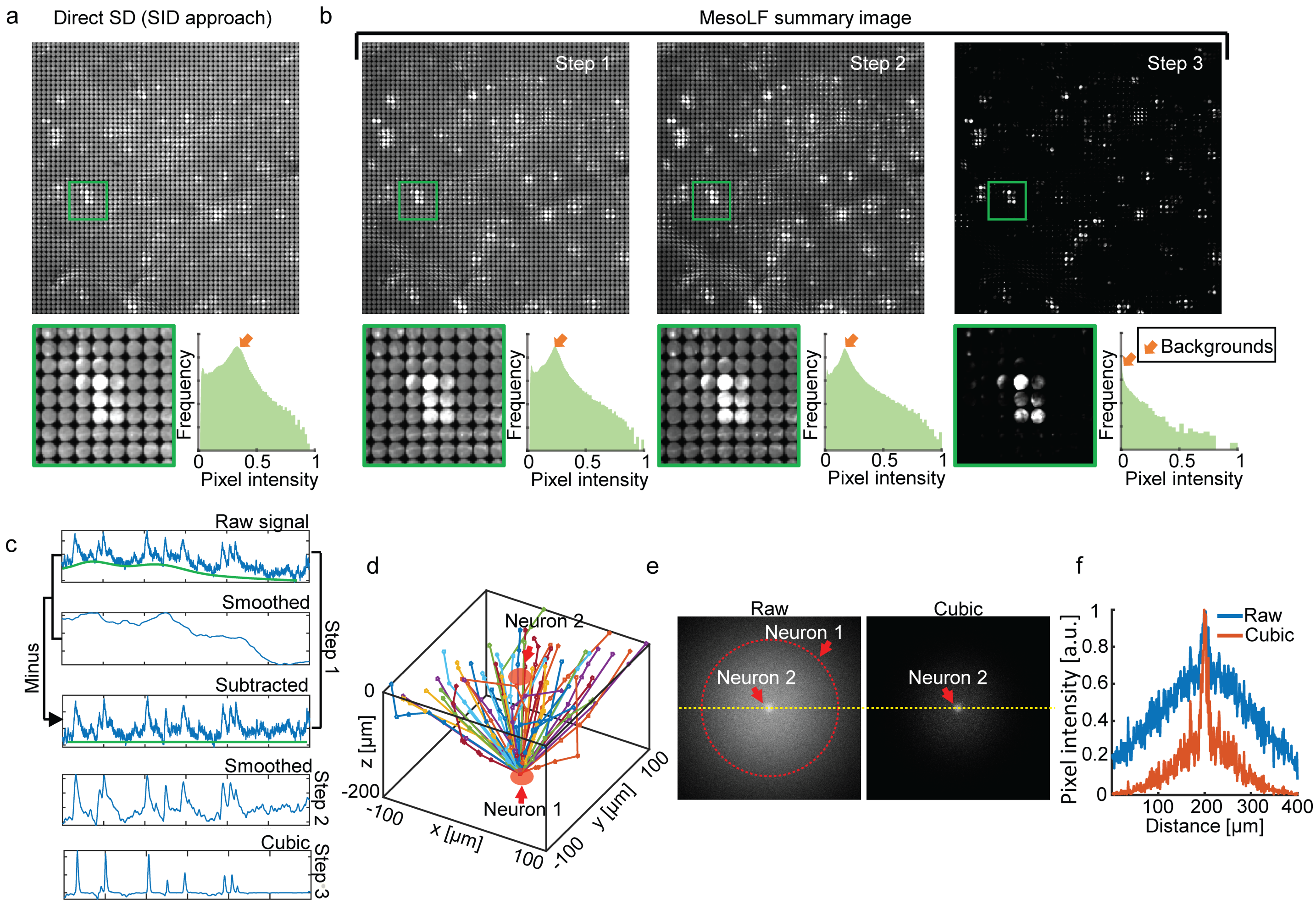


**Supplementary Figure 3**

**Temporal summary image generation.**

(a) Top panel: Standard deviation (SD) image along time calculated directly from uncorrected raw data. Bottom left panel: Zoom into area indicated by green box. Bottom right panel: Intensity histogram of SD image shown in top panel. Orange arrow indicates the mean value of the background in the SD image.

(b) Illustration of the effect of the three preprocessing steps used in MesoLF onto the standard deviation image: In the first step, a large-window smoothed version of the data is subtracted from the raw data to flatten the baselines. In the second step, a small-window smoothing operation is applied in addition, in order to reduce high-frequency noise in the SD image. In the third step, the smoothed data is taken to the third power to enhance contrast. Panels in bottom row are analogous to bottom panels in (a).

(c) Illustration of the effects of the three preprocessing steps in the MesoLF pipeline onto the timeseries of a single pixel in a calcium activity recording

(d) Examples of simulated photon trajectories obtained from Monte-Carlo simulation of two neurons separated by 200 µm along the z axis. The focal plane for data shown in (e)-(f) is chosen to be the x-y plane containing neuron 2.

(e) Intensity in focal plane obtained from Monte-Carlo simulation as in (e). Left panel: raw intensity. Right panel: Third power of left panel

(f) Intensity profile along the dashed yellow line in the two panels shown in (f).

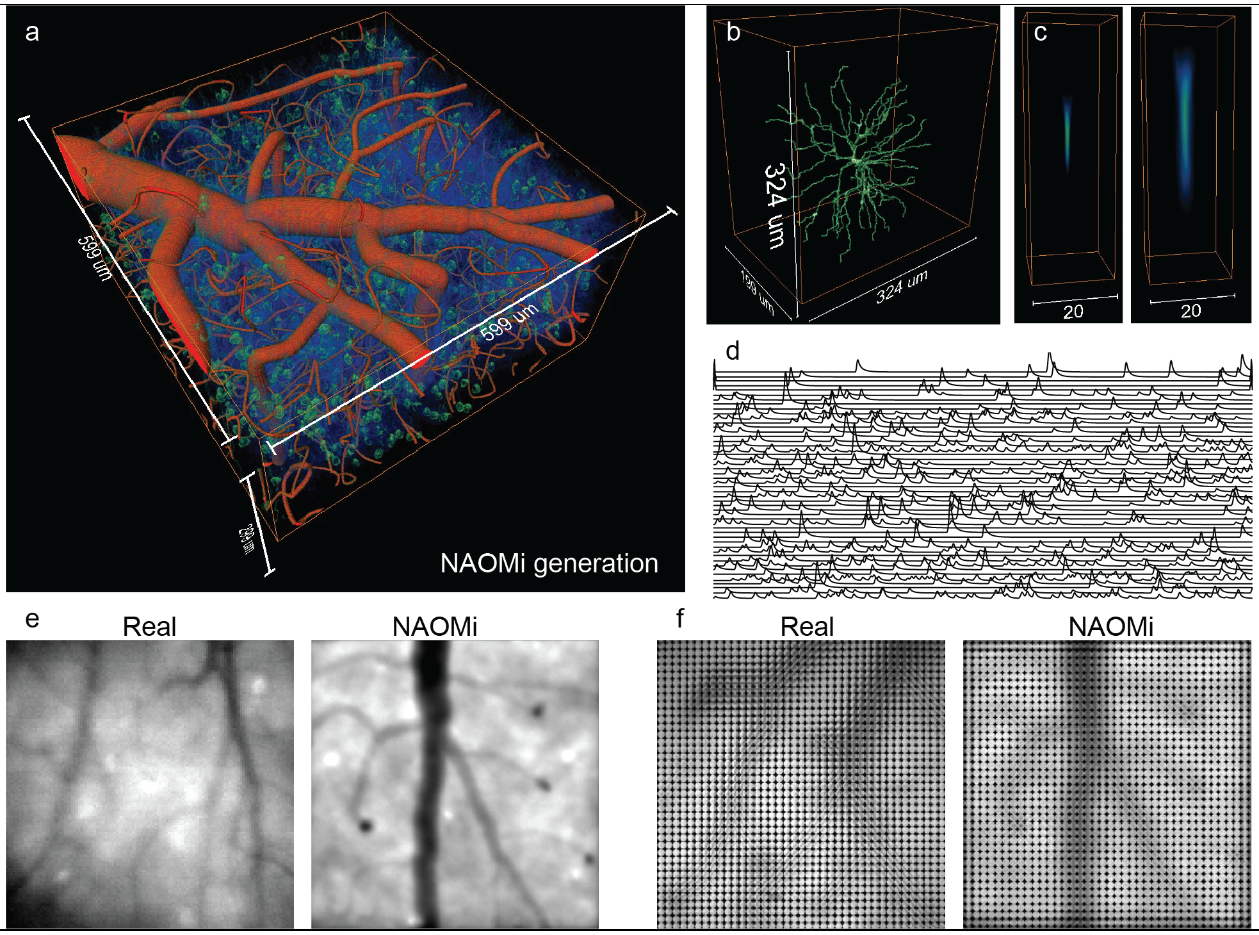


**Supplementary Figure 4**

**Synthetic brain volume and simulation of MesoLF raw data.**

(a) Simulated volume of brain tissue generated using the NAOMi package, filled with blood vessels (red), neurons (green), and dendrites and axons (blue).

(b) An example of a NAOMi-generated neuron with dendrites.

(c) 3D rendering of the optical PSF in free space (left) and in the simulated brain tissue (right).

(d) Simulated calcium activities (fluorescence change normalized to baseline) for neurons.

(e) Experimentally captured wide-field image of brain tissue with GCaMP labeled neurons (left) versus simulated wide-field capture (right). Scale bar: 100 μm.

(f) Experimentally captured light-field image of brain tissue with GCaMP labeled neurons (left) versus simulated light-field capture (right). Scale bar: 100 μm.

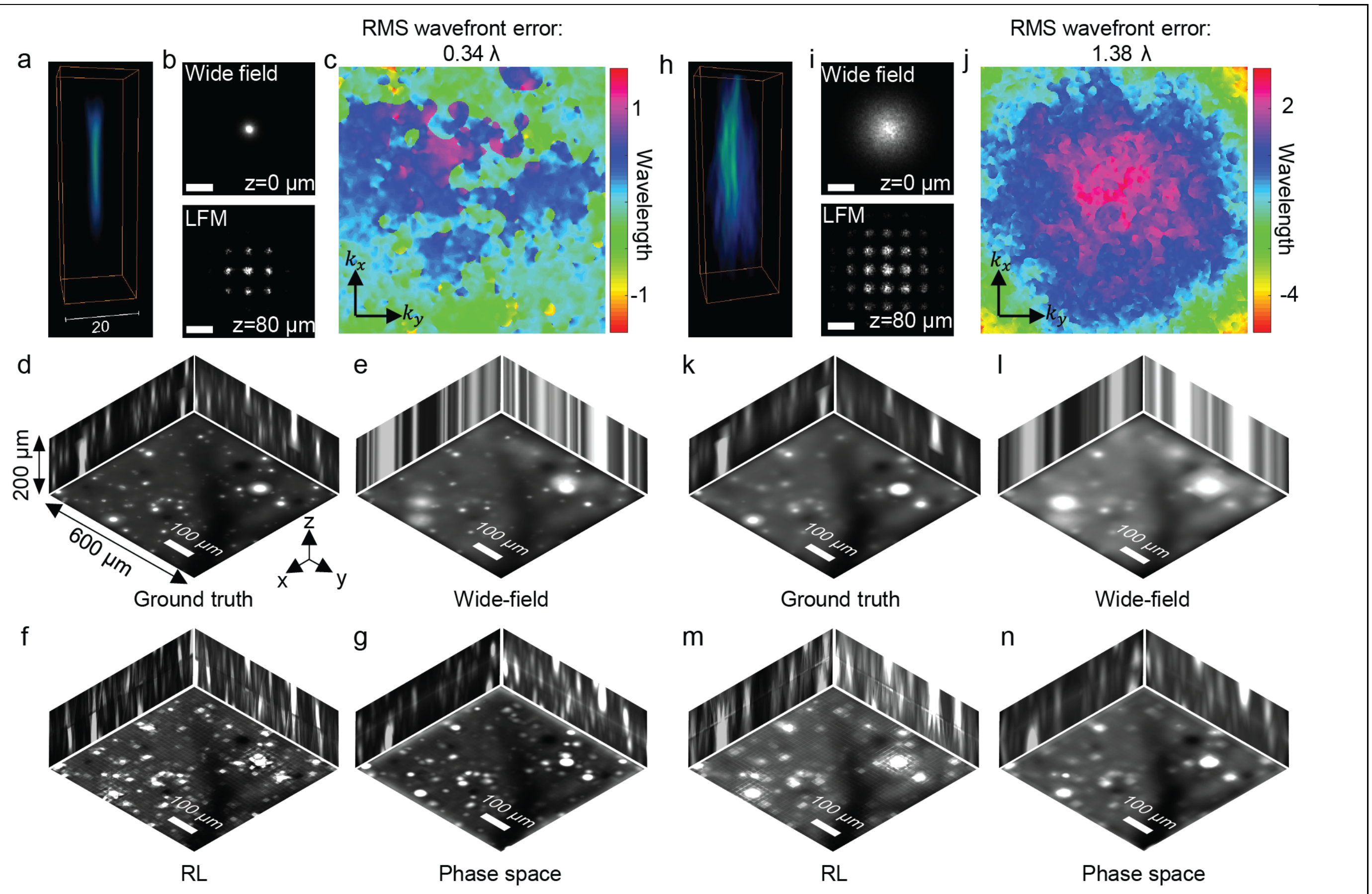


**Supplementary Figure 5**

**Performance comparison of LFM reconstruction methods in the presence of scattering.**

(a) 3D rendering of aberrated, simulated wide-field PSF in the weak scattering scenario.

(b) Intensity cross-section through waist of PSF shown in (a) for wide-field (top, z = 0 µm) and LFM detection (bottom, z = 80 µm).

(c) Wavefront in the pupil plane obtained by backpropagating the PSF shown in (a) to the objective pupil plane.

(d) Maximum intensity projections (MIPs) along three orthogonal axes of the "ground truth" volume obtained by convolving simulated brain tissue volume with simulated PSF shown in (a).

(e) MIPs of simulated volume representing focal stack of wide-field captures.

(f) MIPs of volume obtained by pixel-space Richardson-Lucy (RL) reconstruction of LFM raw data simulated using the PSF shown in (a).

(g) MIPs of volume obtained by MesoLF phase space reconstruction of LFM raw data simulated using the PSF shown in (a).

(h-n) analogous to (a-g) but for the stronger scattering conditions illustrated in (h)-(j).

Scale bars: (b) and (i): 20 µm, (d-g, k-n): 100 µm.

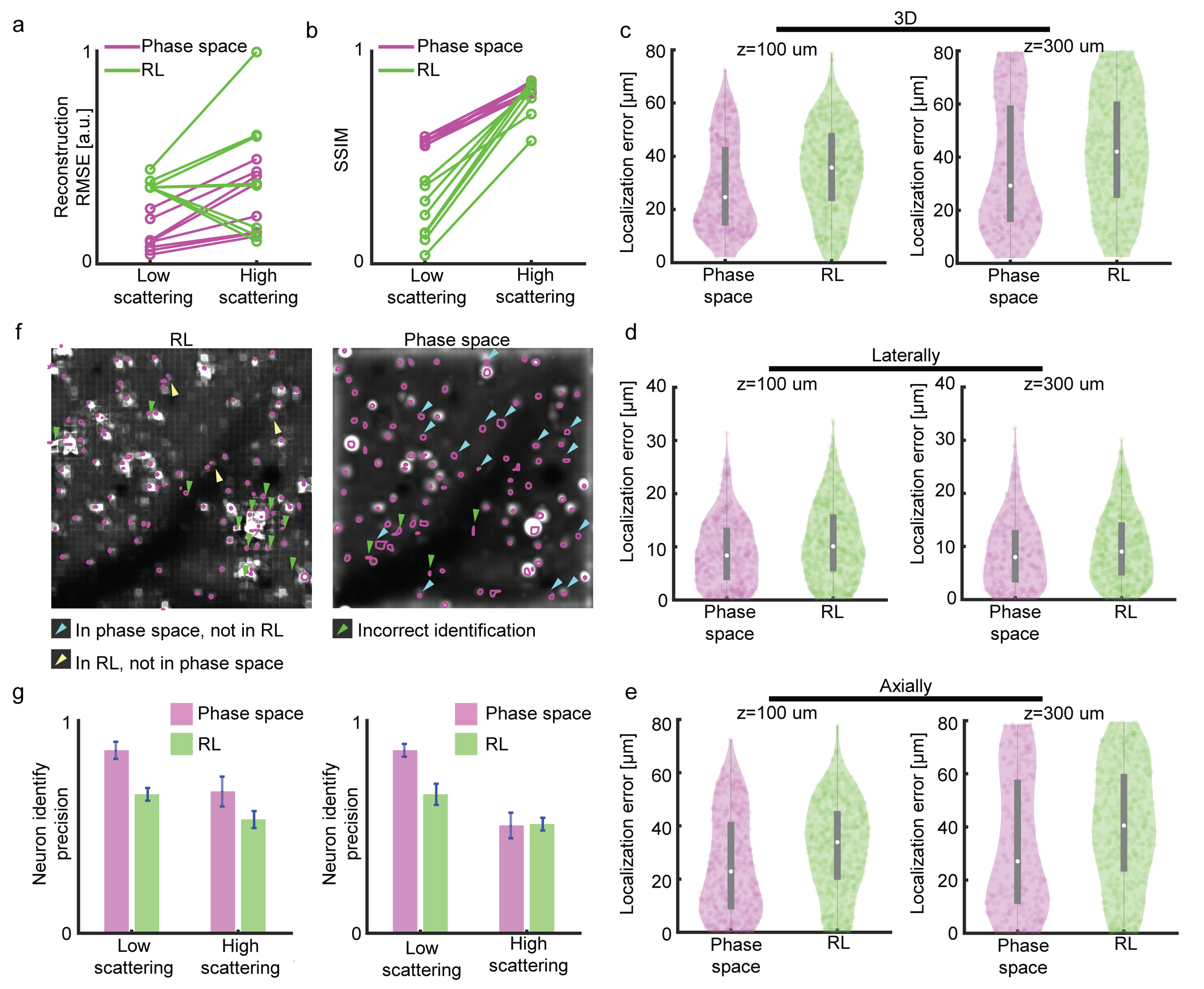


**Supplementary Figure 6**

**Statistical comparison of pixel-space Richardson-Lucy and MesoLF phase space reconstructions.**

(a) Comparison of root-mean-square error (RMSE) between ground truth and reconstructions obtained using pixel-space Richardson-Lucy (RL) reconstruction and MesoLF phase space reconstructions, for the low- and high scattering scenarios. Data points represent runs with different simulated raw data.

(b) Comparison of the structure similarity index (SSIM) values between ground truth and reconstructions, same underlying data as in (a)

(c) Violin plots of the 3D localization errors between ground truth neurons and neurons extracted from phase space (magenta) and RL (green) reconstructions, in the low and high scattering scenarios (depths 100 μm and 300 μm, respectively).

(d) As in (c), but lateral localization error only.

(e) As in (c), but axial localization errors only.

(f) Example plane from reconstructions obtained using RL (left panel) and MesoLF phase space reconstruction (right panel). Magenta circles in the left panel show the segmentation result obtained using the segmentation approach in the SID package (Nöbauer et al., Nat. Methods 14, 2017). Magenta circles in the right panel are segments obtained using MesoLF segmentation. Blue arrows mark the true neurons that appear only in the phase space reconstruction. Yellow arrows mark the true neurons that only appear in the RL reconstruction. Green arrows mark false neuron segments found in reconstructions using both methods.

(g) Comparison of segmentation precision and sensitivity values obtained using phase space (magenta) and RL (green) reconstructions, for the low and high scattering scenarios.

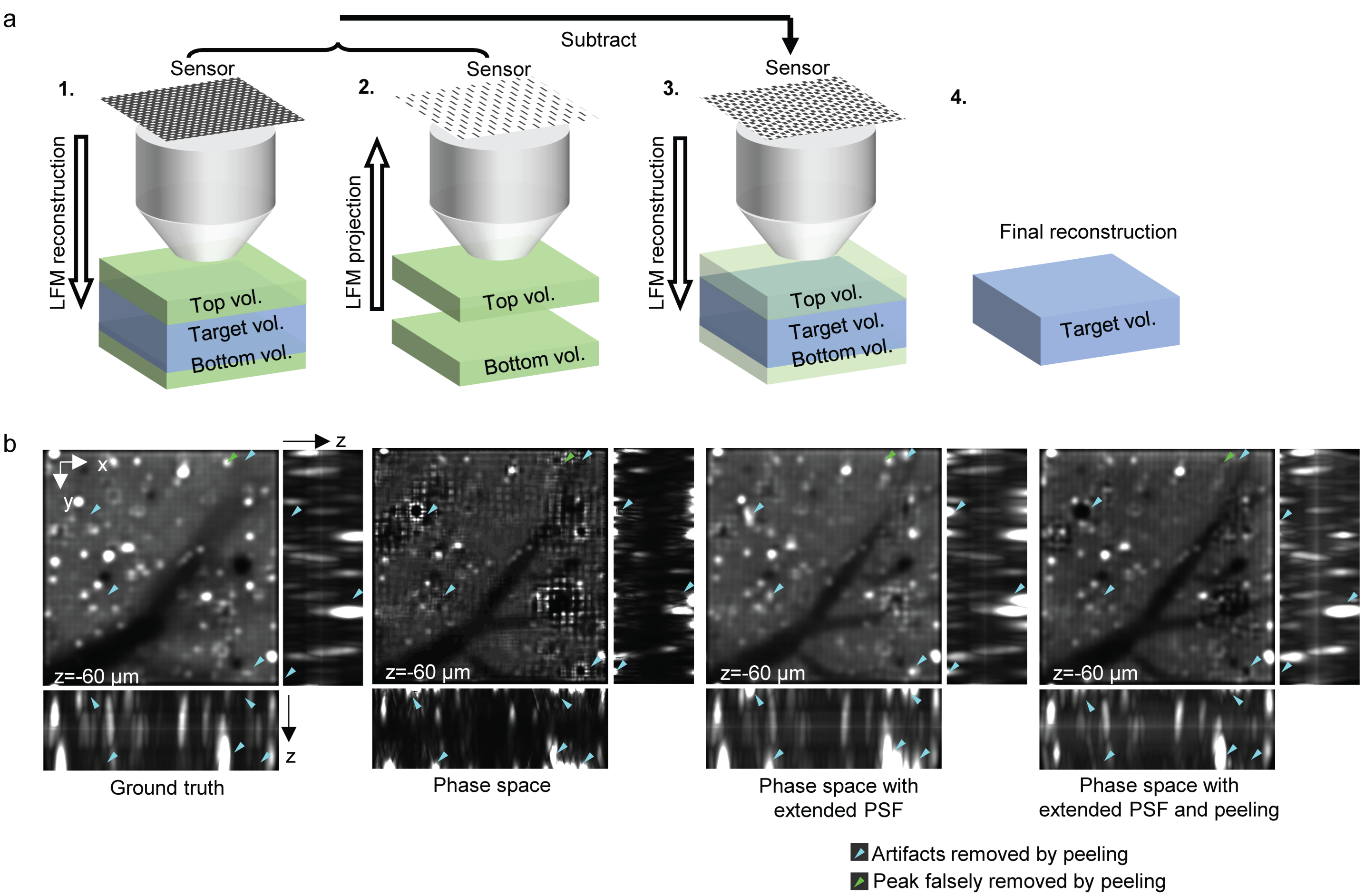


**Supplementary Figure 7**

**Principle of MesoLF reconstruction with background peeling.**

(a) Illustration of the working principle of the MesoLF background peeling technique. First, a full reconstruction is performed using a PSF with an axial extend larger than the targeted reconstruction depth range. Then, the top and bottom layers of the reconstructed volume are numerically propagated to the sensor plane. After subtracting this forward projection from the original raw image, a second run of reconstruction is carried out. Finally, the background layers above and below the target volume are discarded.

(b) Comparison of simulated ground truth and volumetric reconstructions obtained using phase space reconstruction with non-extended PSF, phase space reconstruction with extended PSF, and MesoLF phase space reconstruction with extended PSF and background peeling. The large (x-y) panels are single depth slices at z = -60 µm, whereas the smaller (x-z and y-z) panels are maximum intensity projections. Blue arrows mark bright areas that result from inappropriate reconstructions and artefacts by techniques that do not use background peeling. Green arrows mark bright regions (neurons) that were missed by the background peeling method.

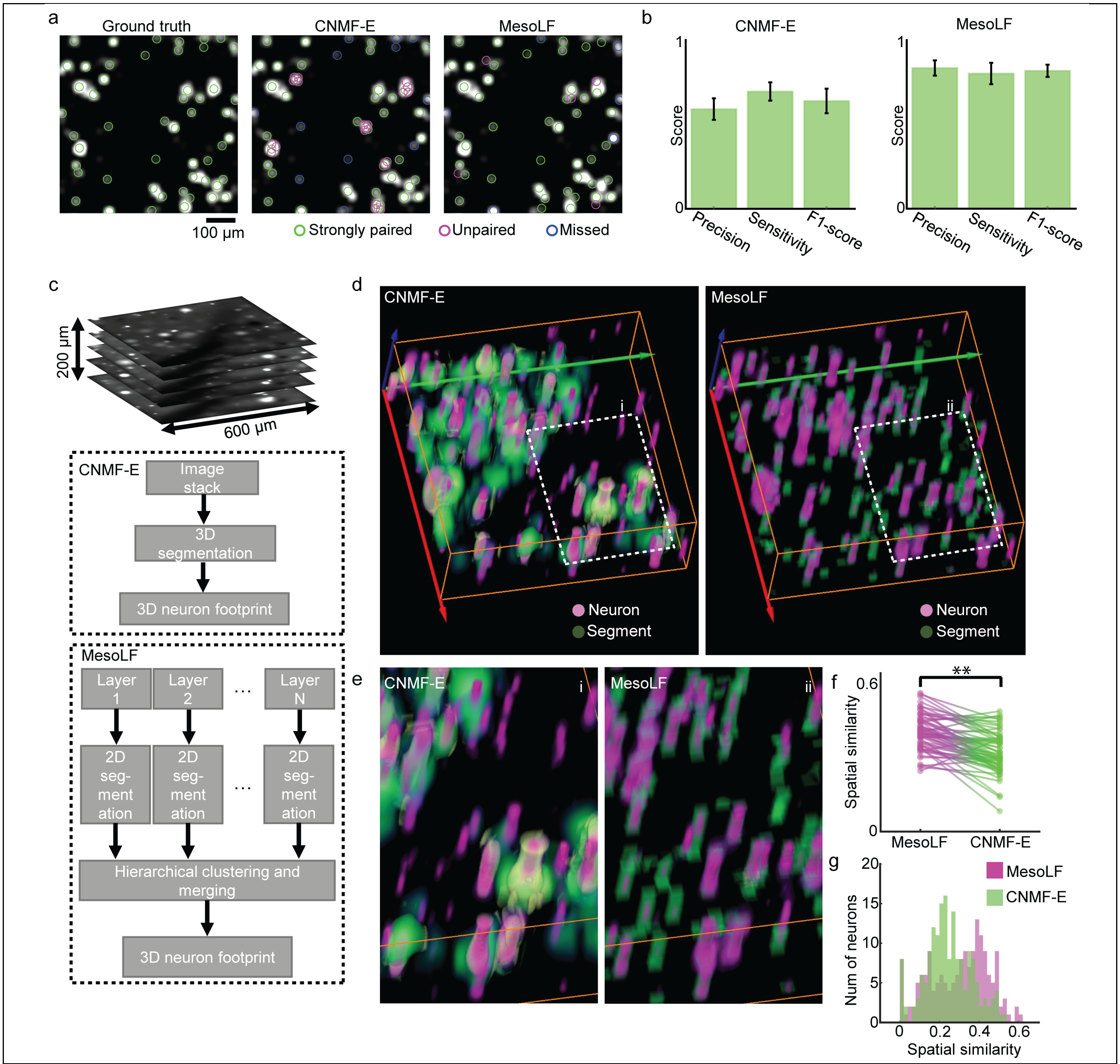


**Supplementary Figure 8**

**Neuron segmentation performance.**

(a) Comparison of segmentation performance of MesoLF versus CNMF-E in a 2D slice from a MesoLF recording in mouse cortex, depth 100 µm. Green circles: segments that strongly match with the ground truth. Blue circles: segments that only appear in the ground truth. Magenta circles: segments that are not consistent with ground truth.

(b) Comparison of precision, sensitivity, and F1-scores for neuron detection performance in CNMF-E (left) and MesoLF (right).

(c) Top panel: Illustration of 3D volume containing neurons and exhibiting scattering, as used for volumetric segmentation comparisons in remainder of figure. Schematic illustration of segmentation pipelines in CNMF-E (middle box) and MesoLF (bottom box).

(d) 3D rendering of segmentation results from CNMF-E (left) and MesoLF (right). Magenta: Ground truth neurons, green: segments.

(e) Zooms into areas indicated by dashed line in (d).

(f) Comparison of the spatial similarity index of neurons paired between ground truth and CNMF-E versus MesoLF.

(g) Histogram of spatial similarity indices of segmented neurons compared to ground truth by both methods (same data as in (f)).

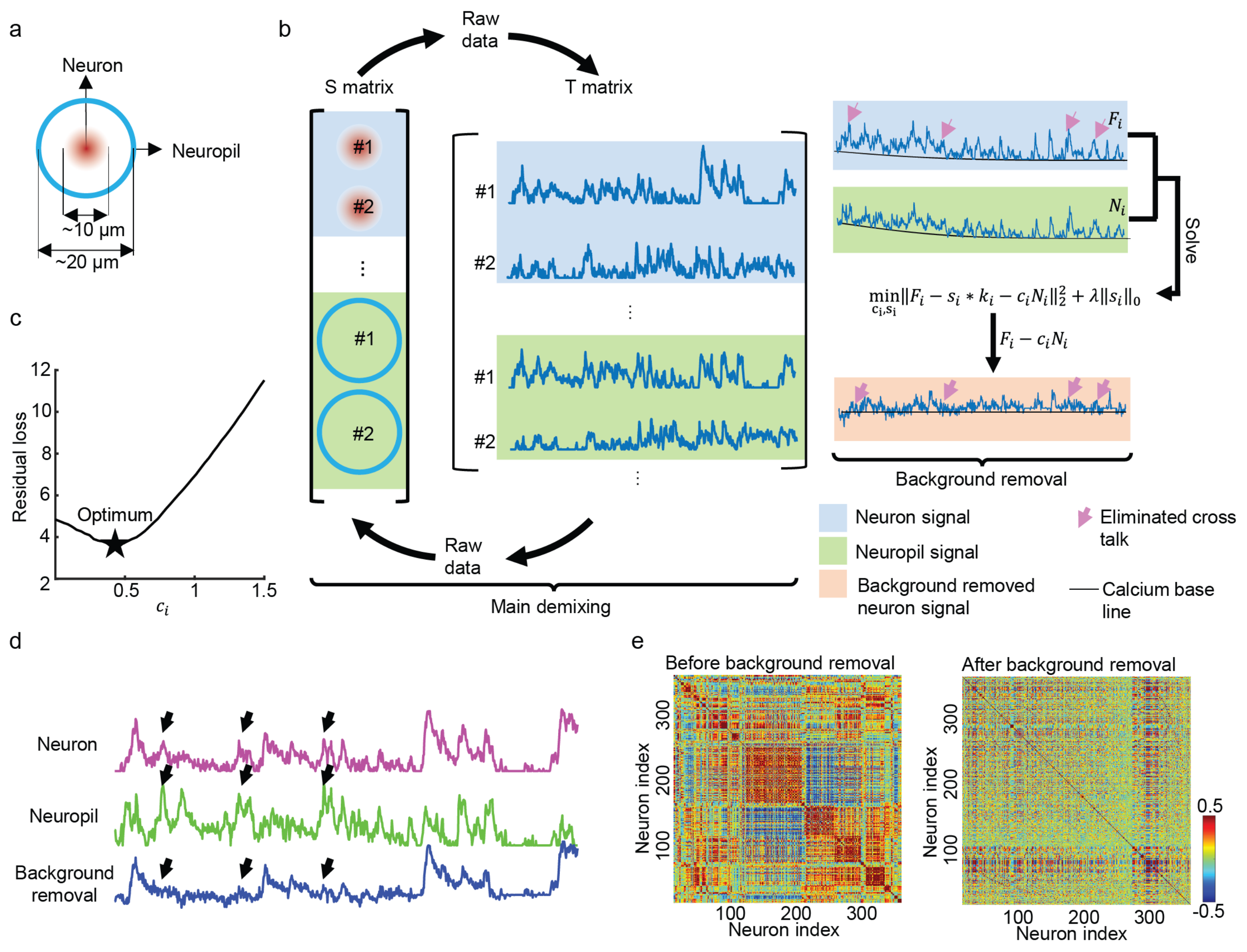


**Supplementary Figure 9**

**Core-shell demixing.**

(a) Illustration of core-shell model used for demixing neuronal signals from local background.

(b) Illustration of the MesoLF main demixing stage, during which the spatial footprints and temporal traces of both core (neuron) and shell (background, neuropil) components are updated alternatingly and iteratively. Rightmost panels: After the main demixing step, a second optimization is run to infer the amplitude mixing coefficient, which subsequently allows to subtract shell contaminations from the core region.

(c) Illustration of loss as a function of a $c_i$ coefficient value during the greedy search to solve demixing subproblem. Star indicates minimum of loss.

(d) Example traces of a core (neuron) trace, the associated shell (background, neuropil) trace, and the background-removed trace. Black arrows indicate peaks in the core trace that result from crosstalk from the shell signal and get removed by the demixing procedure.

(e) Matrix of Pearson correlation coefficients between all pairs of neuron activity traces without background subtraction (left) and with background subtraction (right), for an example dataset.

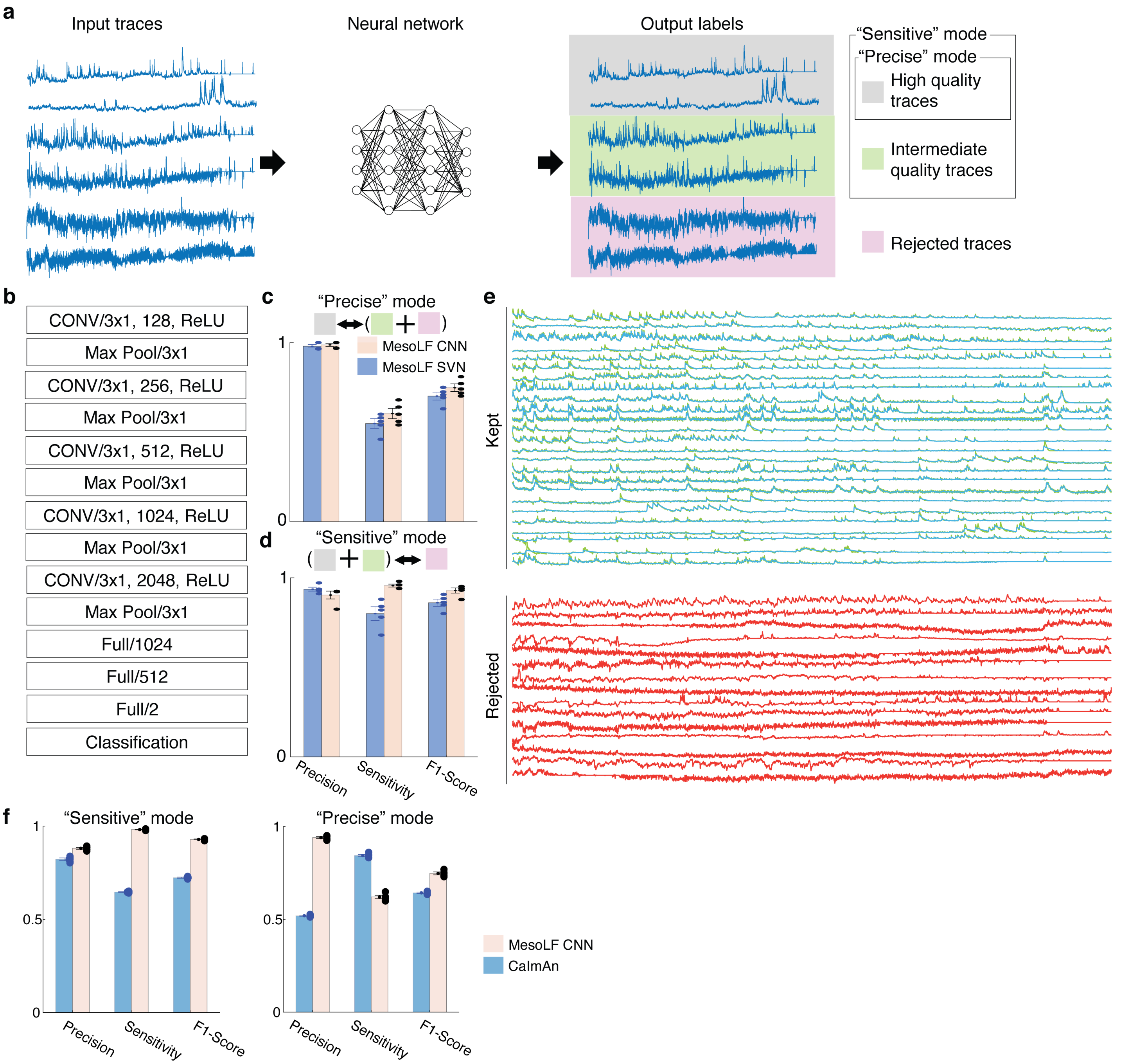


**Supplementary Figure 10**

**Trace classification using supervised learning.**

(a) Schematic illustration of the MesoLF trace classification method. Raw traces are classified by a neural network based on their signal to noise ratio (SNR) and temporal shape. The network was trained on manually labeled data in two different configurations: In "precise" mode, only high-quality traces are retained. In sensitive mode, both high- and intermediate-quality traces are selected to minimize the number of falsely rejected traces.

(b) Structure of the convolutional neural network used for MesoLF trace classification.

(c) and (d) Precision, sensitivity, and F1-scores achieved by MesoLF neural network classifier (yellow) compared to a Support Vector Machine (blue) trained on the same data, evaluated in n = 5 held-out test datasets.

(e) Examples of traces kept and rejected by the neuronal network.

(f) Comparison of precision, sensitivity and F-score for time trace classification between the MesoLF CNN and the time trace post-selection logic in the CaImAn package (i.e., fit with autoregressive model, estimation of noise level from Fourier transform, estimation of and peak-to-noise ratio (PNR), threshold on PNR), for two different settings, “sensitive” (left panel) and “precise” (right panel). Scores were evaluated on a human-labelled test dataset that had not been seen during training. CaImAn precise vs. sensitive scenario chosen by scanning PNR threshold.

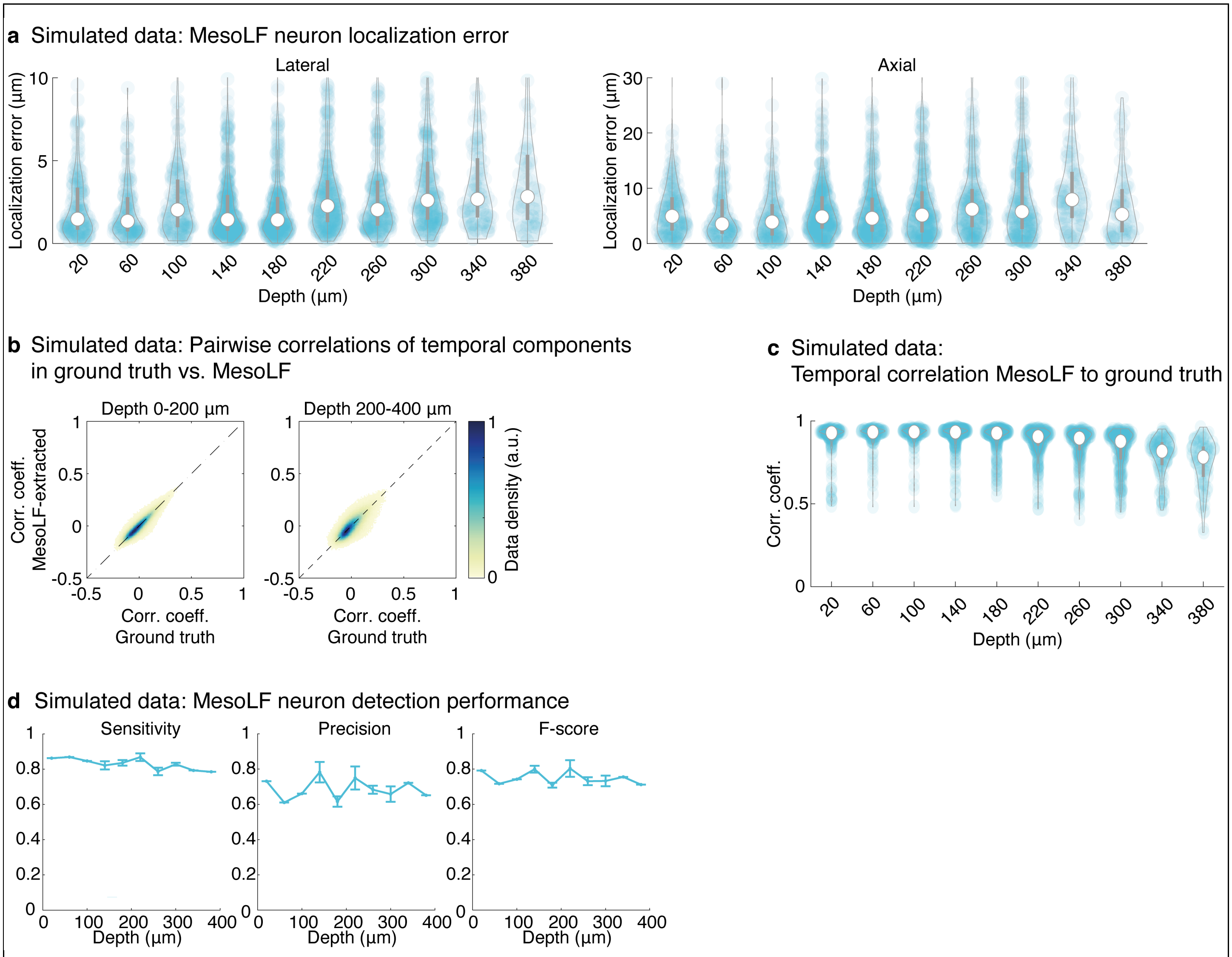


**Supplementary Figure 11**

**Validation of performance of full MesoLF pipeline on simulated data.**

(a) Distributions of lateral (left panel) and axial (right panel) neuron localization error between ground truth and MesoLF-extracted neuron positions from simulated data (with labelled and active neuropil, active neuron density: 14,000 per mm$^3$). White circle: median. Vertical grey bar: Interquartile range. Transparent blue disks: data points. Transparent violin-shaped areas: Kernel density estimate of data distribution

(b) Density plots of pairwise correlations between all pairs of neuronal activity traces in MesoLF, plotted versus the pairwise correlation of the corresponding trace pair in ground truth, for simulated data with labelled and active neuropil. Left panel: Depth range 0-200 µm. Right panel: 200-400 µm. Simulation parameters as in (a)

(c) Distributions of temporal correlation between MesoLF-extracted traces and corresponding traces in ground truth, for simulated data (with labelled and active neuropil), as a function of depth. Violin plot symbols as in **a**. Simulation parameters as in (a)

(d) Neuron detection scores sensitivity, precision and F-score achieved by MesoLF on realistically simulated dataset (without labelled neuropil, active neuron density: 14,000 per mm$^3$) as a function of depth. Error bars: std. err.

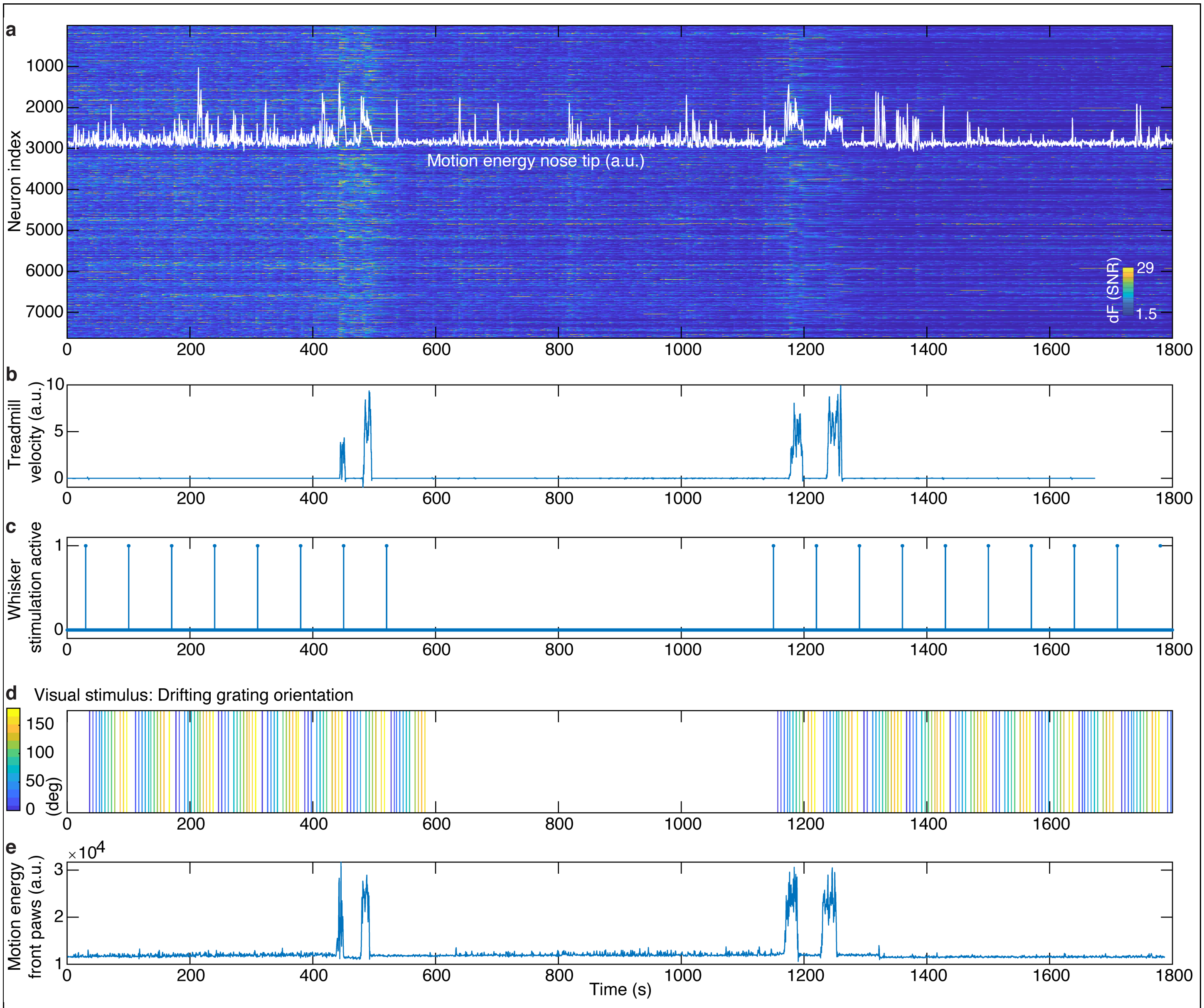


**Supplementary Figure 12**

**30-min MesoLF recording with stimulus and behavior data.**

(a) Background: Heat map of temporal signals extracted from 30-minute MesoLF recording at 18 volumes per second in mouse cortex. Depth range: 100–300 µm. 7651 neurons, sorted by depth (lower neuron index corresponds to lower depth). Overlaid white trace: Motion energy extracted using Facemap Python package (Methods) for region of interest containing nose tip, mouth and base of whiskers. Note the large fraction of global neural activity events that coincide with spontaneous motor behavior of the nose tip, as especially visible during the absence of whisker and visual stimuli in the middle part of the recording.

For same recording as in (a):

(b) Treadmill velocity

(c) Whisker stimulation timepoints (value of 1 indicates whisker stimulation brush is moved in)

(d) Timepoints and orientation of drifting grating visual stimulation

(e) Motion energy extracted using Facemap Python package for region of interest containing front paws (see Methods)
