## Supplementary Notes for "Mesoscale volumetric light field (MesoLF) imaging of neuroactivity across cortical areas at 18 Hz"

### SUPPLEMENTARY NOTE 1: MESOLF OPTICAL DESIGN AND DATA ACQUISITION

The MesoLF optical system (Supplementary Fig. 1a) has been implemented on the widely used commercial "2p-RAM" multiphoton imaging platform[1,2]. In our realization, MesoLF is entirely mounted onto the 2p-RAM vertical breadboard which can be translated along three axes and rotated around one axis via a motorized gantry, although other mounting solutions are possible, including simple static mounting on a fixed breadboard.

The 2p-RAM optical system consists of the excitation laser path, the objective, and the fluorescence detection arm. Of these major parts, MesoLF only uses the 2p-RAM main objective. The original 2p-RAM design[1] also contains a simple, low-resolution wide-field one-photon imaging arm for sample positioning and focusing. The MesoLF excitation and imaging path is mounted in place of this original wide-field arm. Besides its volumetric light field imaging capability, the MesoLF camera signal can also be used directly for sample positioning, thus taking over the function of the original wide-field arm. With the microlens array in place, the camera captures an LFM image, i.e., the image appears segmented into illuminated circles forming behind each microlens. The lateral spatial information that is directly present in this LFM raw image is still sufficient for sample positioning and focusing without any postprocessing, for example, by simply zooming out the camera stream display such that each microlens image circle appears as a single effective pixel to the experimenter. Alternatively, the light field refocusing algorithm[3] can be run in real time to compute at wide-field focal slice from the LFM data.

Fundamentally, LFM trades in lateral resolution for axial resolution to achieve 3D imaging within the constraint of the space-bandwidth product transmitted by the pre-LFM (wide-field) optical system. It is thus important to design a pre-LFM optical system that maximizes the achievable space-bandwidth-product to begin with. We therefore replaced the off-the-shelf back-to-back achromat pair that is used as a simple tube lens without specifically optimized corrections for high-resolution widefield imaging in the original design with a custom-designed tube lens system that corrects aberrations of the main objective, primarily in a 515–535 nm emission range of green (GCaMP) calcium indicators (see further below). The tube lens also provides secondary corrections in the ~600–620 nm emission range of red calcium indicators (jRCaMP, jRGECO[4]).

The MesoLF custom tube lens (together with the 2p-RAM objective) transmits a space-bandwidth product of ~25 Megapixels. This contrasts with ~2–4 Megapixels for a standard LFM microscope, as we demonstrated in previous work. This space-bandwidth-product can be utilized for high-lateral-resolution wide-field imaging (without gaining depth information), or, as in an LFM, be translated to ~20 Megavoxels of volumetric information, with lower lateral resolution but gaining depth information. In MesoLF, we choose the LFM design parameter such that one voxel is on the scale of the size of a single neuron.

MesoLF excitation and fluorescence light are coupled into/out of the 2p-RAM microscope "tube", i.e., the section between the main infrared/visible dichroic beamsplitter that is located just behind the main objective, and the first lens of the 2p-RAM PMT detection module, via a motorized 45-degree fold mirror. The 2p-RAM design already contains a motorized linear stage at this position for switching to the original wide-field arm. We replaced the original rigid mirror mount with an adjustable kinematic mount to allow precise positioning of the mirror at the intersection of the optical axes of objective and our custom-designed tube lens and at a 45-degree angle with respect to both axes. This alignment is achieved by replacing the objective with a cage system containing two iris apertures that define the optical axis, centering a green alignment laser onto that axis, and overlapping the back-reflections from the various surfaces in the tube lens as seen on a card with a hole for the incoming laser beam, held below the fold mirror.

The MesoLF tube lens is the centerpiece of our optical system design: When used without a microlens array, our tube lens upgrades the 2p-RAM to be a high-resolution mesoscopic wide-field microscope with a 10× magnification and diffraction-limited resolution at NA 0.4 across a 4-mm FOV.

The 2p-RAM objective is well-corrected for focusing the excitation laser beam at NA 0.6. In the GCaMP spectral range, however, axial and lateral chromatic aberrations, as well as spherical aberrations and coma, remain uncorrected so that the simulated resolution of the objective alone (assuming a paraxial tube lens) drops to less than ~70 line pairs per millimeter at NA 0.4, for a GCaMP-compatible emission band of 515–535 nm. The 2p-RAM objective is also not corrected for field curvature, with a sag of ~160 µm, curving towards the objective from the center of the 2p-RAM's 5-mm FOV. An additional difficulty arises from the relatively long distance of 210 mm between the objective and the closest position where a tube lens can be positioned, and the limited tube diameter and the size of the fold mirror, which restricts the achievable FOV–NA trade-off.

Our custom tube lens addressing all these issues consists of three cemented doublets in a configuration based on the Petzval objective design form[5]. This lens corrects aberrations to such a degree that diffraction-limited resolution is restored at an NA of 0.4 up to a field radius of ~1.9 mm (RMS wavefront error plot in Supplementary Fig. 1b and ray fan plot in Supplementary Fig. 1c) and reduces field curvature to ~60 µm (sag at center relative to edge of 4-mm FOV). This corresponds to a simulated resolution of ~600 line pairs per millimeter (lp/mm) across the full FOV of 4 mm in diameter (see modulation transfer function (MTF) plot in Supplementary Fig. 1d). We performed the tolerancing of this lens design in Zemax while mechanical design of the housing, further tolerancing, manufacturing, and optical verification was performed by Jenoptik Optical Systems. Anti-reflection coatings were applied to the lens elements to allow for >95% transmission in the wavelength range of 450-670 nm. In the 600–620 nm emission range of red calcium indicators, the tube lens is designed to achieve a resolution of ~500 lp/mm up to a FOV diameter of 2 mm. The outer surface of tube lens mount is threaded (thread size M75.5 × 0.5) to allow mounting into the 2p-RAM housing, replacing the original tube lens.

To hold the dichroic beamsplitter (Semrock FF505-SDi01 short-pass dichroic, 80 × 50 mm) and emission filter (Semrock Brightline FF01-525/39, ⌀2"), we designed a custom mechanical part (Supplementary Fig. 1a, bottom left inset) that slips onto the rear, cylindrical section of the tube lens and is held in place by a metal strap clamp such that emission light is reflected off the dichroic beamsplitter towards the camera. We designed a custom kinematic mount (right inset in Supplementary Fig. 1a) that holds the microlens array (MLA, RPC Photonics MLA-S100-f12, square grid, pitch 100 µm, f = 1.2 mm, F-number 12.5, diced to 42 × 42 mm) and is mounted onto the camera housing. The kinematic mount allows for precise adjustment of the tip/tilt angle, the translation along the optical axis, and the rotation around the optical axis via four fine adjustment screws. The microlens array is aligned with respect to the camera by illuminating it with a 40-mm-diameter collimated green laser beam and adjusting the aforementioned degrees of freedom until uniform and focused spots can be observed behind all microlenses. Rotation is adjusted to line up the microlens grid exactly parallel to the camera pixel grid.

The camera (Teledyne DALSA Falcon 4-CLHS 86M, 86 Megapixels, 6 µm pixel pitch, 12 bit, global shutter, 16 fps full frame rate) is read out via a CameraLink-HS cable and a PCIe frame grabber card (Teledyne DALSA Xtium-CLHS PX8) in the control PC, and image data is streamed to one of two available software-defined RAID-0 arrays of two PCIe flash disks each (2× Samsung 970 EVO 2TB and 2× Sabrent Rocket 2280 4TB, respectively) using a custom data acquisition application written in VisualC# .NET. The magnified image covers an area of ⌀40 mm on the camera, which corresponds to ~7000 × 7000 pixels. This subset of pixels can be read out at 18 fps, resulting in a raw data rate of ~1320 MB/s.

The MesoLF illumination path was designed and optimized in Zemax (non-sequential mode) to provide uniform illumination intensity across the full 4-mm FOV. It consists of a mounted blue LED (Thorlabs M470L3, 470 nm center wavelength, 650 mW), an adjustable asphere collimator (Thorlabs SM2F32-A), an iris aperture for adjusting illumination NA, excitation filter (Chroma ET470/40x, ⌀2"), an engineered diffuser for creating a flat-top intensity profile (RPC Photonics EDC-10-15027-A 2S, 2" square), relay lens (Edmund 45-418, f=300, ⌀3"), and three fold mirrors. The relay lens is positioned at a distance equal to its focal length from the diffuser and the same distance from the illumination-side rear image plane of the tube lens. This arrangement provides telecentric, homogeneous illumination both in the image plane and the focal plane in the sample. We routinely image with an illumination power of ~15 mW post-objective, which corresponds to a power density of ~1.2 mW/mm$^2$ in the sample, a value comparable to our previous LFM imaging methods and typical wide-field imaging.

We measured the as-built wide-field resolution before installing the microlens array and with the camera positioned in the rear image plane of the tube lens by imaging a high-resolution USAF target (Supplementary Fig. 1e). On the USAF target, we obtain 20% MTF contrast for group 8-5, which corresponds to 406 lp/mm. This measured resolution is lower than the design resolution primarily because in the realized microscope, the bandwidth of the emission filter is 39 nm rather than the 20 nm for which the tube lens achieves diffraction-limited resolution. We chose to use this wider filter to maximize signal at higher frame rates at a certain expense of resolution. After adding and aligning the MLA and moving the camera-MLA assembly backwards using a precision linear stage such that the MLA was positioned in the rear image plane, we proceeded to characterizing the resolution achievable through LFM imaging and reconstruction. For an USAF target positioned at z = +90 µm from the objective's native focal plane, we find an MTF contrast of ~50% for group 6-5, which corresponds to 102 lp/mm, thus achieving the design goal of about cell-resolving lateral resolution. To estimate the axial resolution, we LFM-imaged ⌀6 µm fluorescent beads and take their FWHM reconstructed axial extent as a proxy for the resolution. For beads positioned at z = -30 µm relative to the native focal plane, we find an axial FWHM of ~32 µm. The reconstructed lateral FWHM of the 6 µm beads is ~7 µm, compatible with the lateral resolution measurement obtained from imaging an USAF target. We note that – as demonstrated in detail in previous publications[6,7] – the neuron discriminability in the presence of scattering is predominantly determined by the temporal dynamics of neuroactivity, specifically the difference in neuronal activity between neighboring neurons, and to a lesser extent, by optical resolution.

### SUPPLEMENTARY NOTE 2: MOTION CORRECTION

When imaging in awake, behaving animals, rigid motion correction of functional imaging data is usually necessary, even at small fields of view. At the mesoscopic field of view that MesoLF is capable of capturing, mere rigid motion correction becomes insufficient: At this scale, we routinely observe non-rigid deformations on the scale similar to the size of a neuron (see main text).

A variety of methods have been described for correcting brain motion such that it does not disturb the extraction of calcium activity[8–10]. However, none of these methods are suitable for LFM imaging since they require the ability to shift the acquired raw image laterally. In LFM, the raw image does not directly consist in an image of the sample but requires computational reconstruction to yield an estimate of the sample volume. In the raw image, lateral and angular information contained in the light field emanating from the sample are folded into each other and, in addition, light reaching the sensor is masked by the apertures of the microlenses (Supplementary Fig. 2a-b). In principle, it would be conceivable to reconstruct the individual LFM raw frames one-by-one and then apply existing motion correction, but this would be excessively costly computationally and break the design strategy underlying MesoLF. Methods for excluding motion-affected frames[6] or interpolating neuron activities from specific kernels[6] only approximate the original neuronal activities during motion events. As part of the MesoLF pipeline, we developed a new method for correcting motion of brain images in an LFM detection geometry without any frame-by-frame reconstructions.

Our method starts by rearranging the pixels in the raw image into a 2D array of so-called sub-aperture images. A sub-aperture image consists of all the sensor pixels that have the same position relative to their nearest microlens. If there are $M \times M$ microlenses and the image formed by each is sampled using $N \times N$ sensor pixels, then the raw image (which has dimensions $M \times M \times N \times N$) can be rearranged into $N \times N$ sub-aperture images with dimensions $M \times M$ each. Each sub-aperture image corresponds to a perspective view onto the sample from a specific angle. We now perform motion registration on the central sub-aperture image, which consists in the central pixel behind each microlens. We use the FFT-based registration pipeline NoRMCorre[10] to obtain the non-rigid motion information, which is then applied inversely to all the sub-aperture images.

Subsequently, we re-arrange them into the original LFM raw image pixel arrangement (Supplementary Fig. 2c).

To verify the validity of this approach, we plot the correlation matrix between the shift vectors computed individually for each sub-aperture image (i, j) in Supplementary Fig. 2b: The motion shifts imparted onto the sub-aperture images are all highly correlated with each other, except for those at the edges of the correlation matrix. These sub-aperture images at the edges correspond to angular perspectives from large angles, and hence they do not contain a large amount of information on lateral shifts. The observation that all sub-aperture motion vectors are highly correlated validates our strategy to apply the shifts calculated from the central sub-aperture image to all others.

This approach has several advantages: First, it acts on sub-aperture images which can be built directly from the raw images without prior reconstruction. Second, the sub-aperture images reproduce continuous sample features as continuous image features, compared to the periodically folded structure of LFM data in the raw images. This allows the use of established motion correction algorithms to extract the motion shifts. Third, the lower number of pixels in the sub-aperture image makes the registration almost a hundred times faster compared to using whole raw images. Fourth, since we apply motion correction individually to small patches of the full field of view, complex non-rigid deformation and motion of the brain tissue can be corrected.

After motion correction, the correlation between subsequent frames of the central sub-aperture images is increased compared to the raw data (Supplementary Fig. 2d). MesoLF corrects the motion of individual smaller patches of the full field of view in parallel. Whereas the motion vectors found within one patch (typically ~700 × 700 µm as shown in the illustration of the motion correction pipeline in Supplementary Fig. 2c) are only slightly different, the motion vectors seen for different tiles across the full 4-mm FOV differ significantly (Supplementary Fig. 2f), which underscores the necessity of non-rigid correction. This assertion is supported further by the observation that in this mesoscale FOV regime, frame-to-frame correlations remain low when using global rigid motion correction only, but improve markedly when correcting different tiles individually, as is done in the MesoLF pipeline (Supplementary Fig. 2g).

### SUPPLEMENTARY NOTE 3: SUMMARY IMAGE GENERATION

After background subtraction, motion correction, and filtering, the MesoLF pipeline generates a temporal summary image which is subsequently used to find the initial neuron positions. In this temporal summary image, pixels with strongly varying activity over time stand out, whereas weakly varying or entirely static background is de-emphasized. Simply taking the standard deviation of each pixel along the time axis can achieve such an effect. However, neurons can be surrounded by temporally active neuropil, and light scattering in brain tissue will blur the images of the neuron cell bodies and mix the ballistic cell body images with scattered photons, which may originate from other neurons. Under these conditions, directly taking the standard deviation to generate a summary image results in suboptimal contrast (Supplementary Fig. 3a).

To improve the contrast of the temporal summary image, we pre-process the data in three steps prior to taking the standard deviation: First, we perform pixel-wise high-pass filtering to eliminate slow background. This is achieved by subtracting a low-pass-filtered version of each trace (moving average with large window length) from itself. As shown in Supplementary Fig. 3c, this results in a flat baseline, so that after taking the standard deviation along time, the contrast generated by the actual neuronal activity is improved compared to taking the standard deviation without prior high-pass filtering (left panel of Supplementary Fig. 3b compared to Supplementary Fig. 3a).

As a second preprocessing step, to further distinguish active neurons from high-frequency background noise, we apply a temporal filter with a small convolutional window size (Supplementary Fig. 3c and middle panel of Supplementary Fig. 3b).

To investigate the spread of scattered photons around a ballistic neuron image, we conducted Monte Carlo simulations of two neurons separated axially by 200 µm in brain tissue (Henyey-Greenstein model, scattering length 60-80 µm, anisotropy factor g=0.9) (Supplementary Fig. 3d). We found that in the raw camera image, scattered photons from the deeper neuron (neuron 1) add a broad halo around the image of the in-focus neuron (neuron 2), causing a blurred background. To mitigate this effect, we apply a third preprocessing step in which we take the raw images to the third power, which increases local contrast and makes in-focus neurons more recognizable (Supplementary Fig. 3e-f). Since the subsequent neuron segmentation relies on local contrast only, this non-linear rescaling does not negatively affect neuron segmentation performance.

### SUPPLEMENTARY NOTE 4: PHASE SPACE DECONVOLUTION WITH BACKGROUND PEELING

A key factor in achieving the neuron detection performance in scattering media demonstrated by our approach is our novel MesoLF deconvolution algorithm, which we also refer to as artefact-free phase space reconstruction with background peeling. Our solution builds on the approach introduced by Lu et al.[11] and is related to techniques used in ptychography to reconstruct an image from projections of the sample along a set of angles.

Before introducing the MesoLF phase space reconstruction algorithm, let us briefly restate the conventional Richardson-Lucy LFM reconstruction approach[12,13] which may be loosely characterized as operating in “camera pixel space” as opposed to “phase space”. The difference will be clarified in what follows.

#### “Pixel space” Richardson-Lucy reconstruction

A common implementation of a light-field microscope consists in a wide-field microscope with a microlens array (MLA) placed in the native imaging plane. We denote the sample space coordinates as $(x_1, x_2, z)$ and sensor plane coordinates as $(s_1, s_2)$. The point spread function (PSF) of such an LFM can be written as follows[12]:

$$h(x_1, x_2, z, s_1, s_2) = \left|\mathcal{F}_{f_\mu}\{U(x_1, x_2, z, s_1, s_2)\Phi(s_1, s_2)\}\right|^2. \quad (1)$$

Here, $\mathcal{F}_{f_\mu}\{\cdot\}$ denotes the operator in Fresnel approximation that propagates a field by a distance $f_\mu$ along the optical axis. $U(x_1, x_2, z, s_1, s_2)$ is the field in the native image plane generated by a point source at $(x_1, x_2, z)$, given by[14]

$$U(x_1,x_2,z,s_1,s_2)=\frac{M}{f_{obj}^2\lambda^2}\exp\left(-\frac{iu}{4\sin^2(\alpha/2)}\right)\int_0^\alpha P(\theta)\exp\left(-\frac{iu\sin^2(\theta/2)}{2\sin^2(\alpha/2)}\right)J_0\left(\frac{\sin(\theta)}{\sin(\alpha)}v\right)\sin(\theta)\mathrm{d}\theta,$$
$$v\approx k\sqrt{(x_1-s_1)^2+(x_2-s_2)^2}\sin(\alpha),$$
$$u\approx 4kz\sin^2(\alpha/2)\,, \tag{2}$$

with parameters defined as in Ref. 12.

In Eq. 1, $\Phi(s_1,s_2)$ is the phase shift function describing an MLA, parametrized by microlens pitch $d$ and focal length $f_\mu$

$$\Phi(s_1,s_2)=\iint \mathrm{rect}\left(\frac{t_1}{d}\right)\mathrm{rect}\left(\frac{t_2}{d}\right)\exp\left(-\frac{ik}{2f_\mu}(t_1^2+t_2^2)\right)\cdot$$
$$\mathrm{comb}\left(\frac{s_1-t_1}{d}\right)\mathrm{comb}\left(\frac{s_2-t_2}{d}\right)\,\mathrm{d}t_1\mathrm{d}t_2. \tag{3}$$

To reconstruct the 3D sample from the captured image, the continuous sample and sensor space is discretized[12]. LFM can then be modelled as a linear system $H$ which maps the 3D sample space onto 2D sensor space:

$$\sum_{x_1,x_2,z} H(x_1,x_2,z,s_1,s_2)X(x_1,x_2,z)=Y(s_1,s_2). \tag{4}$$

Here, $Y$ is the discrete sensor image and $X$ is the 3D brightness distribution of the sample. The weight matrix $H$ can be computed from Eq. 1, which describes how photons emitted from the voxel $(x_1,x_2,z)$ contribute to the signal received at pixel $(s_1,s_2)$. The weight matrix $H$ can be simplified by exploiting the translational periodicity of the MLA, which implies

$$H(x_1,x_2,z,s_1,s_2)=H(x_1+D,x_2+D,z,s_1+D,s_2+D), \tag{5}$$

where $D$ is the pitch of microlens array in units of the sensor pixel pitch. Note that the sample coordinates $(x_1,x_2)$ can be rewritten as

$$x_1 \coloneqq (m_1-1)D+u_1,$$
$$x_2 \coloneqq (m_1-1)D+u_1. \tag{6}$$

We denote the system response from the sample area in front of the central microlens with coordinates $(m_{10},m_{20})$, projected in the sample space as

$$H_{FOR}(u_1,u_2,z,s_1,s_2)\coloneqq H((m_{10}-1)D+u_1,(m_{20}-1)D+u_2,z,s_1,s_2), \tag{7}$$

The overall forward projection matrix $H$ from sample to sensor can then be written as

$$H(x_1,x_2,z,s_1,s_2)\coloneqq\ H_{FOR}(u_1,u_2,z,s_1-(m_1-m_{10})D,s_2-(m_2-m_{20})D). \tag{8}$$

Correspondingly, the forward model can be written as

$$Y(s_1,s_2)=\sum_z\sum_{u_1,u_2}\sum_{m_1,m_2} H_{FOR}(u_1,u_2,z,s_1-(m_1-m_{10})D,s_2-(m_2-m_{20})D)X(m_1,u_1,m_2,u_2,z). \tag{9}$$

For simplicity, we use the notation $\mathbf{H}_{\mathrm{for}}(X)=Y$ to represent the forward projection in LFM defined by Eq. 9.

Similarly, for backward projection from the sensor to the sample, we introduce the new coordinates $n$ and $v$:

$$s_1\coloneqq(n_1-1)D+v_1$$

$$s_2 := (n_2 - 1)D + v_2 \quad (10)$$

The inverse response of the sensor area behind the central microlens $(n_{10}, n_{20})$ in sensor space can then be written as

$$H_{BACK}(x_1, x_2, z, v_1, v_2) := H(x_1, x_2, z, (n_{10} - 1)D + v_1, (m_{20} - 1)D + v_2) \quad (11)$$

The transmission matrix $H$ can then be represented using $H_{BACK}$ as

$$H(x_1, x_2, z, s_1, s_2) = H_{BACK}(x_1 - (n_1 - n_{10})D, x_2 - (n_2 - n_{20})D, z, v_1, v_2) \quad (12)$$

Thus, the backward model is

$$X(x_1, x_2, z) = \sum_{v_1, v_2} \sum_{n_1, n_2} H_{BACK}(x_1 - (n_1 - n_{10})D, x_2 - (n_2 - n_{20})D, z, v_1, v_2) Y(n_1, v_1, n_2, v_2) \quad (13)$$

which we write as a shorthand using the notation $\mathbf{H}_{\text{back}}(Y) = X$. It is common to use the well-known Richardson-Lucy (RL) algorithm to iteratively update $X$ from $Y$ and $H$. In each iteration, RL computes the new estimate $\hat{X}^{(t)}$ from the result of the previous iteration, $\hat{X}^{(t-1)}$, via

$$\hat{X}^{(t)} \leftarrow \hat{X}^{(t-1)} \odot \mathbf{H}_{\text{back}} \left( \frac{Y}{\mathbf{H}_{\text{for}}\left(\hat{X}^{(t-1)}\right)} \right), \quad (14)$$

where $\odot$ denotes element-wise multiplication.

#### Phase space deconvolution

Before we introduce the innovations in the MesoLF reconstruction algorithm, it is necessary to outline the phase space reconstruction technique proposed by Lu et al.[11]. We start by expanding $H_{FOR}$ into the sensor plane coordinates that are based on the periodicity of microlens array.

$$H_{FOR}(u_1, u_2, z, n_1, v_1, n_2, v_2) = H((m_{10} - 1)D + u_1, (m_{20} - 1)D + u_1, z, (n_1 - 1)D + v_1, (n_2 - 1)D + v_2) \quad (15)$$

Similarly, by denoting $Y_{v_1,v_2}(n_1, n_2)$ as $Y((n_1 - 1)D + v_1, (n_2 - 1)D + v_2)$, the LFM forward projection operation can be written as

$$Y_{v_1,v_2}(n_1, n_2) = \sum_{z} \sum_{u_1,u_2} \sum_{m_1,m_2} H_{FOR}(u_1, u_2, z, n_1 - (m_1 - m_{10}), v_1, n_2 - (m_2 - m_{20}), v_2)\ X(m_1, u_1, m_2, u_2, z) \quad (16)$$

Here $Y_{v_1,v_2}(n_1, n_2)$ is called the phase space response of LFM under the phase space coordinate $(v_1, v_2)$. For simplicity, we denote

$$H_{PFOR,v_1,v_2,z}(u_1, u_2, n_1, n_2) = H_{FOR}(-u_1, -u_2, z, n_1, v_1, n_2, v_2) \quad (17)$$

and

$$X_z(u_1, u_2, m_1, m_2) = X(m_1, u_1, m_2, u_2, z). \quad (18)$$

Then,

$$Y_{v_1,v_2}(n_1,n_2) = \sum_z \sum_{u_1,u_2} \sum_{m_1,m_2} H_{PFOR,v_1,v_2}(0-u_1, 0-u_2, n_1-(m_1-m_{10}), n_2-(m_2-m_{20}), z) X_z(u_1,u_2,m_1,m_2)$$
$$= \sum_z \left(H_{PFOR,v_1,v_2} * X_z\right)|_{(0,0,:,:)} \quad (19)$$

Here, $*$ denotes convolution between $(u_1, u_2)$ and $(m_1, m_2)$. Note that the convolution between $H_{PFOR,v_1,v_2}$and $X_z$ results in a 4D array, so we need to select a 2D slice from the 4D array to match the dimension of $Y_{v_1,v_2}(n_1, n_2)$.

For backward projection, recall that the traditional LFM backward projection is (Eq. 13):

$$X(x_1, x_2, z) = \sum_{v_1,v_2} \sum_{n_1,n_2} H_{BACK,v_1,v_2}(x_1-(n_1-n_{10})D, x_2-(n_2-n_{20})D, z) Y_{v_1,v_2}(n_1,n_2) \quad (20)$$

This formulation already uses what we now recognize as the sensor data in phase space representation $Y_{v_1,v_2}(n_1, n_2)$, thus the LFM backward projection in phase space simply remains the same as in the "pixel space" formulation:

$$X_{v_1,v_2}(x_1, x_2, z) = \sum_{n_1,n_2} H_{BACK,v_1,v_2}(x_1-(n_1-n_{10})D, x_2-(n_2-n_{20})D, z) Y_{v_1,v_2}(n_1,n_2) \quad (21)$$

For simplicity, let us write the phase space forward projection operation as $\mathbf{H}_{FOR}^{v_1,v_2}(\cdot)$, and the backward propagation as $\mathbf{H}_{BACK}^{v_1,v_2}(\cdot)$. With an initialization value for the sample space estimate $X_{init}$ and a captured image $Y$, the phase space LFM reconstruction algorithm is

$$\hat{X}^{(t)} \leftarrow \hat{X}^{(t-1)} + \alpha_{t-1}\hat{X}^{(t-1)} \odot \mathbf{H}_{BACK}^{v_1,v_2}\left(\frac{Y}{\mathbf{H}_{FOR}^{v_1,v_2}\left(\hat{X}^{(t-1)}\right)}\right). \quad (22)$$

The update rule Eq. 22 runs over all phase space coordinates $(v_1, v_2)$. The advantage of phase space reconstruction comes from the freedom to choose the order of updating the $(v_1, v_2)$ slices. By choosing to update first those slices that correspond to high spatial frequencies, the reconstruction can be accelerated compared to pixel-space Richardsion-Lucy[11]. The update weight $\alpha_{t-1}$ can also be chosen, with larger $\alpha_t$ usually leading to a sparser result.

**Algorithm 1. MesoLF artifact-free phase space reconstruction**

Initialize $X^{(0)} = X_{\text{init}}, t = 0$.

Rearrange captured image $Y$ into phase space image $Y_{v_1,v_2}$.

Calculate the anti-aliasing filter radius $\omega(z)$ for the different $z$ depths.

**while** $t$ < T do:

$t = t + 1$

**for** $(v_1, v_2)$ in phase space, do:

$$X^{(t)} = X^{(t-1)} + \hat{X}^{(t-1)} + \alpha_{t-1}\hat{X}^{(t-1)} \odot \mathbf{H}_{BACK}^{v_1,v_2}\left(\frac{Y}{\mathbf{H}_{FOR}^{v_1,v_2}(\hat{X}^{(t-1)})}\right)$$

**for** z in all depths, do:

$$X_z^{(t)} = X_z^{(t)} * e^{-\frac{(x^2+y^2)}{2\left(\frac{\omega(z)}{2\sqrt{2\log 2}}\right)^2}}$$

**end**

**end**

**end**

**return** $X^{(t)}$

#### MesoLF artefact-free phase space reconstruction (without peeling)

The phase space update rule given in Eq. 22 is computationally more efficient than the established "pixel-space" Richardson-Lucy approach but still suffers from similar reconstruction artifacts, which originate from aliasing due to the depth-dependent sampling density in sample space in an LFM geometry[12]. In our MesoLF reconstruction approach, we therefore combine phase space reconstruction with the use of different filters for different depth slices of the sample volume[15]. Specifically, following each update step, the slice at depth $z$ of the sample space estimate is convolved with a Gaussian filter. The FWHM $\omega(z)$ of the filter is given by

$$\omega(z) = \min\left(\left|\frac{d_{sens}^{mla}}{d_{tl}^{mla}}\right|\frac{z''}{d_{tl}^{mla} - z''}\left|D - \frac{D}{2z'''\left|z''' - d_{sens}^{mla}\right|}\right|, \frac{D}{2}\right), \tag{23}$$

with

$$z'' = M^2(z - f_{obj}) + d_{tl}^{mla},$$

$$z''' = \frac{d_{sens}^{mla}(d_{tl}^{mla} - z'')}{d_{tl}^{mla} - z'' - d_{sens}^{mla}}$$

Here, $d_{sens}^{mla}$ is the distance between the MLA and the sensor, $d_{tl}^{mla}$ is the distance between tubelens and MLA. $\omega(z)$ reaches its maximal size in the native image plane to mitigate the large square-shaped artefacts that otherwise appear there. $\omega(z)$ decreases with increasing distance from the native image plane. The full MesoLF artefact-free reconstruction algorithm (without peeling) is listed as pseudocode in the box entitled Algorithm 1.

**MesoLF phase space reconstruction with background peeling**

In one-photon imaging, it is very difficult to confine the axial extent of the illumination. This has the consequence that tissue above and below the depth range targeted for LFM reconstruction will also be excited and emit fluorescence. Such out-of-range signals mix with in-range signals, which degrades the reconstruction contrast and reduces the signal-to-background ratio. Furthermore, blood vessels in superficial tissue layers expand and contract periodically and thus generate a temporally modulated shadowing effect for fluorescence emanating from further below. While blood vessels also absorb at the excitation wavelength, we expect the effect of excitation shadowing to be partially compensated by scattering and ultimately diffusion of illumination light, so that excitation shadowing effects are much less pronounced than emission shadowing. These temporal modulations of emission intensity at blood vessel boundaries manifest themselves in the temporal activity summary images that are generated during MesoLF processing. If left untreated, these spurious activity signals would lead to false neuron candidates because some sections of these vasculature-induced activity patterns would appear in the output of a naïve segmentation algorithm as neuron-like segments. To solve this problem, we developed a technique which we refer to as "background peeling" and which can more faithfully extract true signals from background-contaminated data by iteratively reconstructing and removing out-of-range signals. We describe this approach in what follows.

We start by explicitly including out-of-range regions above and below the targeted reconstruction range into the LFM forward model:

$$Y = HX + H^A X^A + H^B X^B = [H^A, H, H^B][X^A; X; X^B] \quad (24)$$

where $H$ is the forward projection matrix for the target depth range, $X^A, X^B$ are the defocused out-of-range volumes above and below the target range, and $H^A, H^B$ are the forward projection matrices for these two volumes. The depth range of $X^A$, $X$, and $X^B$ is denoted as $1, \dots, d_T$, $d_{T+1}, \dots, d_{T+S}$, and $d_{T+S+1}, \dots d_{T+S+B}$. The difficulty in solving this problem stems from the unknown $X^A$, $X^B$, and the unknown range of $H^A$,$H^B$. These large uncertainties render potential solutions to Eq. 24 non-unique.

To approach this challenge, we use a ballistic PSF $H'$ with a larger $z$ depth range to perform reconstruction, i.e., $col(H') > col(H)$ where $col(\cdot)$ returns the column number. The extended PSF $H'$ serves as an approximation to the true underlying $[H^A, H, H^B]$, which is the true forward projection matrix of the imaging system in the presence of scattering but remains unknown. After reconstruction, only depths $d_{T+1}, \dots, d_{T+S}$ are retained as the reconstruction of the target depth range. Reconstruction with such an extended PSF will remove some of the background from out-of-range layers.

On the other hand, since the scalar products $\langle H_i, H_j \rangle \neq 0$, $\langle H_i^A, H_j \rangle \neq 0$, and $\langle H_i^B, H_j \rangle \neq 0$, reconstructions for depth layer $i$ will exhibit crosstalk with the other layers $j$. In other words, even after the volumes above and below the target range have been removed from the extended-depth PSF reconstruction, they will still have effects on the retained depth layers. Besides, the top and bottom volumes $X^A$ and $X^B$ are often much brighter than the target volume $X$ due to the periodic dilation of the blood vessels and large defocused bottom volume. Therefore, their shadows disturb the reconstructions

**Algorithm 2. MesoLF reconstruction with background peeling**

Captured LFM image $Y$. Initialize $X^{(0)} = X_{\text{init}}$. Hyperparameters $\beta_1, \beta_2$.Prepare the extended PSF $\mathbf{H}'$.

► Run reconstruction (using phase space or pixel space Richardson-Lucy method) using $\mathbf{H}'$ and $Y$, which results in volume estimate $X'$

► Take the top layer of $X'$ as $X_1'$, project it to the sensor plane through $p_1 = \mathbf{H}_{\text{for}}\big(X_1' \cdot (X_1' > \beta_1)\big)$.

► Update captured image $Y' = Y - \beta_2 p_1$.

► Take the bottom layer of $X'$ as $X_D'$, project it to the sensor plane through $p_{T+S+B} = \mathbf{H}_{\text{for}}\big(X_D' \cdot (X_D' > \beta_1)\big)$.

► Update captured image $Y'' = Y' - \beta_2 p_{T+S+B}$.

► Run reconstruction (with phase space or RL method) using $\mathbf{H}'$ and $Y''$, and output $X''$.

► Crop the $X'$ along the z-axis by discarding slices $1, \dots T$ and $T + S + 1, \dots, T + S + B$. The result is $X_{out}$

**Return** $X_{out}$

even with extended PSF reconstruction. We thus developed a "background peeling" technique that removes the forward-projection of the top and bottom layers of $X$ from the sensor plane data after the first reconstruction iteration in order to further reduce crosstalk. The brightest structures in the top and bottom layers will iteratively be removed so that they do not interfere with structures from other depths. A pseudocode listing of MesoLF background peeling technique is given in the box entitled Algorithm 2.

### SUPPLEMENTARY NOTE 5: TISSUE SIMULATIONS AND PERFORMANCE BENCHMARKS ON SIMULATED DATA

To rigorously evaluate and optimize the performance of the MesoLF pipeline in the presence of scattering, in particular with respect to the crucial step of reconstructing the temporal summary image, we performed extensive simulations based on realistic models of cortical tissue that included neurons, neuropil, light scattering, absorption by blood vessels, and LFM image formation.

To synthesize a realistic cortical tissue, we use the Neural Anatomy and Optical Microscopy (NAOMi)[16] package. Using NAOMi, a brain tissue volume is populated with multiple blood vessels, as well as with neuron somata, axons, and dendrites (Supplementary Fig. 4a-b). Neurons and dendrites are assigned synthesized fluorescence activity that reflect their calcium dynamics (Supplementary Fig. 4d).

As light travels through such an inhomogeneous medium, the refractive index differences between cells and other tissue components, as well as absorption by hemoglobin, distort the light wavefront. When propagated through an imaging system, such a distorted wavefront leads to an aberrated PSF (Supplementary Fig. 4c). This PSF can be convolved frame-by-frame with a time series of synthetic sample volumes to obtain a realistic representation of the raw camera data that would be captured when recording calcium activity in vivo.

NAOMi is primarily designed for benchmarking different two-photon imaging technologies as well as calcium imaging analysis algorithms. We therefore adapted the PSF generation code to simulate one-photon wide-field and LFM imaging. We generated a brain tissue volume of 600 × 600 × 300 μm$^3$ in size, where the LFM-targeted depth range corresponds to the central 600 × 600 × 200 μm$^3$ sub-volume, so that both above and below the target range is a 50-μm-thick slice that acts as a defocused, out-of-range background. The labelled out-of-range slices above and below the target range can be much thicker in real experiments in mice, but fluorescence originating from distances much larger than the axial range of LFM leads to a completely blurred background that does not show spatial variation across the FOV of interest and is thus rejected effectively by our initial background rejection steps during preprocessing. It is background from immediately above and below the targeted depth range that most strongly affects signal extraction.

While the average cortical neuron density in the mouse brain, when considered over all cortical layers and across all cortical regions, is reported to be ~85,000–90,000 per mm$^3$, labelling efficiency, driver line selectivity, and lack of neuronal activity during recordings will necessarily lead to lower observed neuron numbers in realistic experiments. Thus, we set the neuron density to 10,000–14,000 neurons per mm$^3$, which we found to lead to synthetic data that is more comparable to the empirically observed values in previous recordings. For the verifications of individual steps of the MesoLF pipeline shown in Fig. 3b-h, neuron density was set to 10,000. As shown in Supplementary Fig. 4e, the simulated one-photon wide-field image (right) closely resembles experimental data (left). Supplementary Fig. 4f demonstrates the same good agreement for simulated and experimental data in the case of LFM imaging. This realistic NAOMi-based simulation of LFM imaging in scattering brain tissue enables us to monitor the performance of the MesoLF calcium extraction algorithms in comparison to an underlying ground truth, which is difficult to access experimentally throughout a comparable depth range and volume size.

Next, we ran both the established pixel-space Richardson-Lucy (RL) as well as our MesoLF phase space reconstruction algorithm on the synthetic LFM datasets and compared the outcomes. First, we run the brain tissue simulation for a shallow depth range (0–200 μm), where the distortion of PSF is relatively small (Supplementary Fig. 5a-b). To quantify the wavefront error, we back-propagate the distorted PSFs to the pupil plane of the objective and compute the RMS wavefront error. In the shallow case, we obtain a value of 0.34 λ compared to the distortion-free wavefront, indicating mild aberration. Even under such a mild scattering, conventional wide-field imaging fails to differentiate signals from different depths (Supplementary Fig. 5e), and Richardson-Lucy reconstructions (Supplementary Fig. 5f) exhibit artifacts along both the lateral and axial directions. Our novel MesoLF phase space reconstruction, on the other hand, results in a visibly better agreement with the ground truth volume, which is here taken as a convolution of the brain tissue with the aberrated PSF, since in the absence of optical aberration correction, this represents the optimally attainable numerical reconstruction of the sample (Supplementary Fig. 5d).

We further compare the reconstruction error under more strongly scattering conditions. The PSF is now more severely distorted (Supplementary Fig. 5j), with an RMS wavefront error of ~1.4 λ (Supplementary Fig. 5j). Both the resulting pixel-space Richardson-Lucy (Supplementary Fig. 5m) and the MesoLF phase space reconstructions (Supplementary Fig. 5n) are blurred due to severe scattering, but artifacts are much more severe in the Richardson-Lucy case, especially near the center of the LFM depth range, which is where the native image plane of the bare microscope is located. In both the low- and high-scattering situations, the proposed MesoLF phase-space reconstruction technique exhibits superior performance compared to the widely used pixel-space Richardson-Lucy algorithm, both in terms of overall reconstruction fidelity and severity of artifacts.

To statistically evaluate these observations, we quantitatively compare the reconstruction methods both regarding the reconstruction error and segmentation bias: The proposed MesoLF phase space method outperforms the established pixel-space Richardson-Lucy method in terms of reconstruction root-mean-square error (RMSE, Supplementary Fig. 6a), both in the low- and high scattering situations. The well-known structure similarity index (SSIM, as implemented by Matlab's ssim() function[17]) also reflects the better performance of MesoLF phase space reconstruction in the case of low scattering (Supplementary Fig. 6b). In the high scattering case, the SSIM values of both methods are high due to the strongly blurred reconstructions, but both variance and mean of the SSIM values obtained for MesoLF phase space reconstructions are smaller, indicating stable performance across different samples.

In the MesoLF pipeline, the main purpose of performing reconstructions is to allow robust and sensitive segmentation of the reconstructed volumes to generate neuron candidates. To quantify the impact of reconstruction performance on segmentation accuracy and precision, we calculated the localization error as the minimum distance between neurons found in reconstructions and neurons in the ground truth dataset. We found that the localization error for neurons segmented from MesoLF phase space reconstructions is smaller than the error resulting from segmenting pixel-space Richardson-Lucy reconstructions, both in the low- and high-scattering scenarios (Supplementary Fig. 6c-e). In particular, the bright and isolated artifacts that pixel-space Richardson-Lucy reconstruction is prone to generate result in many false segments (indicated by green arrows in Supplementary Fig. 6f). The blocky, square artifacts commonly found in Richardson-Lucy LFM reconstructions[12,13] near the native focal plane do not match basic assumptions about neuron shape and are therefore not recognized as neurons by reasonable segmentation approaches. These artifacts are absent in MesoLF phase-space reconstruction, resulting in many additional neuron detections (blue arrows). To further quantify this observation, we classified a segment as a valid neuron if its overlap with the closest ground truth neuron is larger than 50%. As shown in Supplementary Fig. 6g, both the resulting precision score (ratio of number of true positives to sum of true and false positives) and sensitivity score (ratio of number of detected to actual neurons) are higher (better) for MesoLF reconstruction than pixel-space Richardson-Lucy reconstructions.

The comparisons discussed so far in this section do not include the MesoLF background peeling technique described further above (see also schematic in Supplementary Fig. 7a). As shown in Supplementary Fig. 7b, a direct phase-space reconstruction without background peeling and using a simulated PSF that is not extended beyond the targeted 200-µm axial range leads to extremely weak neuron signals at a depth of only 60 µm. This is because both of the out-of-focus tissue slices above and below the target range lead to much brighter contributions than true neuron signals originating from the target volume. The reduced dynamic range of naïve phase space reconstructions leads to many artifacts, and parts of the reconstructed volumes are affected by defocused halos from bright components in top and bottom background volumes (green arrows). When using the same reconstruction algorithm with a PSF with extended axial extent and subsequently discarding the extended depth slices, the signal level at 60 µm depth is markedly improved. However, many bright spots appear that are not true neurons (blue arrows). Such false bright peaks can be observed even more clearly in the x-z and y-z maximum intensity projections. In comparison, our extended PSF MesoLF reconstruction with background peeling can faithfully remove these artifacts and results in reconstructions that show good agreement with the ground truth data in the presence of top and bottom out-of-volume background fluorescence. We compared the neuron localization errors (defined as the overlap ratio between two paired components) resulting from segmentations of reconstructions obtained using these three techniques. The enhanced performance of MesoLF phase space reconstruction with background peeling is apparent both in comparisons of the 3D localization error (Fig. 3d of the main text), as well as the lateral localization error (Fig. 3e of the main text).

In addition to these benchmarks and validations of individual steps of the MesoLF pipeline, we used realistically simulated data (based on NAOMi, as described above) to evaluate the performance of the full MesoLF pipeline. For these studies, the neuron density was increased to 14,000 neurons per $mm^3$ to simulate a more challenging scenario. The performance quantification approach and results are discussed further in the main text, Fig. 4h, Methods, and Supplementary Fig. 11).

### SUPPLEMENTARY NOTE 6: MORPHOLOGICAL SEGMENTATION

To motivate our design of the MesoLF neuron segmentation technique, we start by introducing two established segmentation approaches and subsequently introduce our approach, followed by a comparison.

#### 3D segmentation in CNMF-E

The widely used CNMF-E package[18,8] uses a segmentation approach that is based on convolving a 3D neuron template with the data volume to find candidate neuron positions. While only a crude approximation to the ideal shape of the volumetric image of a neuron, the comparatively low numerical aperture of detection and the scattering nature of sample in the present case justify the use of a simple 3D Gaussian to model the reconstructed shape of a neuron:

$$d = e^{-\frac{x^2+y^2+\alpha z^2}{\sigma_1^2}} \tag{1}$$

Here, $\sigma$ is the expected neuron radius and $\alpha$ is the aspect ratio of the PSF (lateral FWHM over axial FWHM). The first candidate neuron position $(i, j, k)$ is then found by convolving $d$ with the data volume $I_S$ and finding the brightest pixel in the resulting volume:

$$\max_{(i,j,k)} \; I_S * d \tag{2}$$

The 3D shape $d_p = I_S(i,j,k)\, e^{-\frac{(x-i)^2+(y-j)^2+(z-k)^3}{\sigma_1^2}}$ is then considered the first neuron candidate, where $p$ is the candidate index. The shape is subtracted from the data volume, and Eq. 2 is re-evaluated (greedy search). This segmentation method is schematically illustrated in Supplementary Fig. 8c, upper box.

#### Direct 3D segmentation

In reconstructed LFM data, in contrast to two-photon data, some structures in the temporal summary image may have high brightness values but do not correspond to neurons, for example fragments of blood vessels that appear active due to their periodic pulsation. To mitigate this, we attempt to extend the CNMF approach by additionally constructing a Gaussian ring template that represents the dark region immediately surrounding a neuron:

$$d_S = e^{-\frac{x^2+y^2+\alpha z^2}{\sigma_2^2}} - e^{-\frac{x^2+y^2+\alpha z^2}{\sigma_1^2}} \tag{3}$$

Here, $\sigma_2 > \sigma_1$ is the expected radius of the surrounding region. We convolve both $d$ and $d_S$ with the reconstructed summary volume $I_S$, and find the pixel $(i,j)$ which maximizes

$$\max_{(i,j)} \frac{I_S * d|(i,j,k)}{I_S * d_S|(i,j,k)}. \tag{4}$$

We then construct the shape $d_p = I_S(i,j)e^{-\frac{(x-i)^2+(y-j)^2+(z-k)^3}{\sigma_1^2}}$ as the first neuron candidate, where $p$ is the candidate index. Next, we update $I_S$ by subtracting $d_k$, and re-evaluate Eq. 4 to find the next candidate (greedy search).

Despite this core-shell template design, our simulations show that this direct 3D segmentation procedure is not robust enough when it comes to differentiating neurons whose reconstructed volumetric images overlap axially. This is because in such cases these two volumetric images may lead to a misleading brightness peak in the overlap region.

#### 2D segmentations with clustering

To improve segmentation stability, we resort to plane-by-plane 2D segmentation followed by clustering across planes (Supplementary Fig. 8c, lower box). To this end, we assume the target neuron shape to be a 2D Gaussian,

$$d = e^{-\frac{x^2+y^2}{\sigma_1^2}} \tag{5}$$

where $\sigma_1$ is expected neuron radius. In addition, we construct a 2D Gaussian ring shape to represents the dark region surrounding a neuron:

$$d_S = e^{-\frac{x^2+y^2}{\sigma_2^2}} - e^{-\frac{x^2+y^2}{\sigma_1^2}}, \tag{6}$$

where $\sigma_2 > \sigma_1$ is the radius of the surrounding region. We convolve both $d$ and $d_S$ with a given slice $I_S$ of the reconstructed summary volume, and find the pixel $(i,j)$ which maximizes

$$\max_{(i,j)} \frac{I_S * d|(i,j)}{I_S * d_S|(i,j)}. \tag{7}$$

Then, the shape $d_k = I_S(i,j)\, e^{-\frac{(x-i)^2+(y-j)^2}{\sigma_1^2}}$ is considered the first neuron candidate, where $k$ is the candidate index. We then update $I_S$ by subtracting $d_k$, and re-evaluate Eq. 7 (greedy search). Subsequently, we fit an ellipse to the resulting segment and reject candidates not approximated well by an ellipse with aspect ratio <5. This approach allows to robustly reject segments resulting from undesired structures, such as blood vessels.

After plane-wise segmentation, we cluster and merge the 2D segments based on spatial proximity in 3D. The center of merged patches in 3D space is taken as the brightness-weighted center of the voxels in the patch. After merging, a 3D footprint is computed for each of the resulting neuron candidates in the volume, which are used subsequently in the main demixing step of the MesoLF pipeline.

**Comparison of segmentation approaches**

To evaluate the performance of our segmentation method we compare it with a ground truth and a CNMF-E segmentation result as follows: In a single plane from a phase space reconstruction of an LFM raw frame (experimental data from GECI-labelled mouse cortex at a depth of 100 μm), we manually encircle all neurons with ellipses and use the resulting segmentation as the ground truth (Supplementary Fig. 8a). We run our MesoLF segmentation method as well as the CNMF-E segmentation method and compare the results with the ground truth.

As quantified by the well-known precision, sensitivity, and F-scores, our method detects more neurons while missing or incorrectly segmenting fewer (Supplementary Fig. 8b). At the same time, the CNMF-E segmentation method fails to detect neurons that are much dimmer than others and tends to severely over-segment large and bright components. Even though it would be possible to merge some of the over-segmented components later based on their temporal activities, this is undesirable in the context of MesoLF due to the computational overhead incurred by processing superfluous segments.

In a similar manner, we also compared our segmentation method with CNMF-E applied to a 3D dataset. We numerically synthesized a 3D cortical volume of 600 × 600 × 200 $\mu m^3$ with scattering parameters chosen to be comparable to experimental data (Supplementary Fig. 8c, top). A rendering of the 3D segments obtained from CNMF-E and our MesoLF segmentation approach is shown in comparison to the background-free neuronal volume in Supplementary Fig. 8d: CNMF-E results in blurred segments and misses many of the dimmer neurons. Our method uniformly segments both bright and dim neurons, as further highlighted in the zoomed-in views in Supplementary Fig. 8e.

To quantitatively compare segmentation performance for our method in comparison to CNMF-E, we plot the spatial similarity index SSIM of the segments produced by both methods against ground truth in Supplementary Fig. 8f. Our method results in a significantly larger set of similarity indices than CNMF-E. Supplementary Fig. 8g shows histograms of the similarity indices produced by both methods, further demonstrating the improvement achieved by our method.

### SUPPLEMENTARY NOTE 7: DEMIXING AND BACKGROUND REJECTION

Temporal traces extracted from LFM data are prone to be contaminated by signals arising from surrounding neuropil (dendrites and axons). This is because scattering spreads fluorescence photons originating from neuropil into the image of the soma, causing uneven baselines and crosstalk. As part of the MesoLF pipeline, we explicitly take neuropil contamination into account and provide a robust tool to reject these background components and restore low-crosstalk soma signal traces. We model the neuropil in the immediate vicinity of a neuron soma as a spherical shell with an outer diameter that is approximately twice the inner diameter, which is chosen to match a typical neuron size[9,18,19], as illustrated in Supplementary Fig. 9a. We compute both the neuron and neuropil footprints as they would appear on the LFM sensor before the main demixing procedure and use non-negative matrix factorization with sparsity regularization[20,8] to demix the neuron and shell components at the same time. The addition of neuropil "shell" footprints into the main non-negative demixing already reduces neuropil contaminations from the neuron traces, but not completely so: The resulting neuron traces still exhibit uneven baselines (Supplementary Fig. 9b). We therefore introduce an additional optimization step after the main NMF demixing to solve the following problem, which was first proposed for 2pM data in Ref. 9:

$$\min_{\mathrm{c_i,s_i}} \|F_i - s_i * k_i - c_i N_i\|_2^2 + \lambda \|s_i\|_0,$$

where $F_i$ is the activity trace from $i$-th neuron soma, $N_i$ is the trace from the neuropil region surrounding the $i$-th neuron, $s_i$ is a binary time series containing the underlying action potential events and $k_i$ is the GECI response kernel to an action potential. The coefficients $c_i$ are introduced to allow matching the amplitude of neuron and neuropil traces. The above non-convex optimization problem attempts to obtain minimal loss by finding the correct $s_i$ and $c_i$. In contrast to the solution strategy proposed in Ref. 9, we use a greedy search in $c_i$ with a step size of 0.01 and allow values ranging from 0 to 1.5 (Supplementary Fig. 9c), which we find to perform more robustly and efficiently in the MesoLF context. The final output is the cleaned-up neuron trace given by $F_i - c_i N_i$.

Applying this background rejection method results in traces exhibiting fast increase and slow decrease features as would be expected from the known properties of the GECI response kernel. Most false calcium peaks that result from neuropil crosstalk are eliminated (Supplementary Fig. 9d). We compare the Pearson correlation coefficients between all pairs of neuron traces found in a slice at 300 μm depth, with and without our background rejection enabled (Supplementary Fig. 9e): With background removal enabled, the cross-correlation values markedly decrease.

### SUPPLEMENTARY NOTE 8: BLOOD VESSEL REJECTION

As mentioned in several of the preceding sections, the temporal modulation in emission brightness caused by the periodic expansions and contractions of blood vessels has to be carefully mitigated to avoid its false identification as neuronal activity. This is done at several stages throughout the MesoLF pipeline, which we summarize in this section.

First, during preprocessing and summary image generation, the time series of each pixel in the raw camera movie are temporally filtered to exclude brightness variations that are too slow to be compatible with GECI activity.

Second, after summary image reconstruction, we apply the B-COSFIRE filter[21] to highlight blood vessels for subsequent thresholding and masking. For each depth of the reconstructed volume, B-COSFIRE applies a Gaussian filter and a set of shifted Difference-of-Gaussian filters. After a local maximum operation and weighted summation, the output image generated by the filter only contains emphasized versions of the tube-like, elongated shapes associated with blood vessels. We chose B-COSFIRE due to its robustness, high degree of automatization, and reliance of only four parameters for effective operation. Thresholding the B-COSFIRE output results in blood vessel masks. We then exclude the blood vessels from raw reconstructions so that they will not be segmented as neurons.

Third, as mentioned above, after performing segmentation, we fit segments with ellipses and exclude those with an aspect ratio above a threshold.

Finally, we reject any residual vessel-induced temporal signals and associated neuron candidates during the neuron candidate classification stage using supervised machine learning (see below).

### SUPPLEMENTARY NOTE 9: CANDIDATE CLASSIFICATION USING SUPERVISED MACHINE LEARNING

Despite several filtering and demixing stages throughout the MesoLF pipeline, after the final demixing step, a minor fraction of neuron candidate traces still exhibit features incompatible with GECI activity or are otherwise of low quality (low SNR). Therefore, after subtracting the neuropil background from the neuron candidates traces as described in the previous section, we classify traces based on their temporal shape and noise level. We assign all temporal traces to one of three categories: Traces with low noise and few artifacts (high quality), traces with some noise and artifacts (intermediate quality), and traces with high noise and large artifacts (false positives). To accommodate experimenter preferences, we provide two pre-trained classifiers, one optimized to be "sensitive", i.e., to keep traces with both high and intermediate quality, and one optimized to be "precise", i.e., to keep only traces with high quality scores.

To facilitate supervised training of the classifiers, we manually labelled 5,454 traces using a custom GUI tool that allows high-throughput trace annotation in an interactive manner. We use this annotated dataset to train a neural network with 5 convolutional (CONV) layers and three fully connected (FULL) layers (Supplementary Fig. 10b). The input of the network is a vector with 6,000 elements representing the calcium dynamics, downsampled to 6 Hz (for GCaMP6s). We trained the neural network with the Deep Learning Toolbox in MATLAB R2020a with the built-in Adam optimizer for 100 epochs. A customized function was used to monitor the training process and stop the training when accuracy did not improve anymore. For comparison, we also trained a support vector machine (SVM). Both models were tested on held-out test data that had not been seen during training. We designed the test dataset to be balanced, i.e., to contain the same number of traces for each of the classes. As shown in Supplementary Fig. 10b-c, in both "sensitive" and "precise" modes, the proposed neural network approach outperforms the SVM model, both in terms of precision, sensitivity, and F1 scores. In "precise" mode, in which only high-quality traces are kept, both approaches have very high precision and relatively low sensitivity. This indicates that all kept traces actually have high quality while some actual high-quality traces are falsely rejected. This is because the distinction between high- and intermediate-quality traces is not as well-defined as the distinction between true positive and false positive (non-neuron) traces. In sensitive mode, the neural network achieves precision, sensitivity, and F1 scores exceeding 0.9, indicating good performance. We suggest using sensitive mode by default in most applications. In cases where only high-SNR traces are of relevance, the "precise" approach may be more appropriate.

### SUPPLEMENTARY NOTE 10: LIGHT FIELD SOURCE PLANE SHIFTING BASED ON REFOCUSING FOR VOLUMETRIC FUNCTIONAL VERIFICATION DATASET ASSEMBLY

As described in Methods, in the LFM raw data recorded by splitting the fluorescence emanating from a 2pM-excited plane in the sample between a PMT and an LFM detection arm, fluorescence appears to be emanating from an axial plane offset to a relative depth of $\Delta z_0 = -40$ µm from the axial center of the LFM volumetric field of view (the axial center is also known as the LFM native focal plane). This offset is fixed and set by displacing the LFM camera and microlens array backwards from the rear focal plane of the microscope. In what follows we describe how this fixed axial offset is converted to a set of adjacent offsets by computationally shifting the source plane in the light field data.

The absolute depth of the excited plane was set by translating the microscope objective into and out of the sample, thus shifting the 2pM focal plane and the center of the LFM volume together to a series of absolute depths $z$ below the dura mater, maintaining the -40 µm offset. This results in a series of planar 2pM movies, which are further analyzed using the CaImAn pipeline (Methods), and a corresponding series of LFM raw data movies, an individual frame of which we denote as $I_{z,\Delta z_0}$. Each such raw LFM image $I_{z,\Delta z_0}$ encodes light emanating from the 2pM focal depth $z$, but at a relative shift to the LFM native focal plane at $\Delta z_0$. In other words, if these LFM images $I_{z,\Delta z_0}$ were 3D-reconstructed, the result would be volumetric movies in which all fluorescence would appear to be emanating from a single quasi-planar region offset by -40 µm from the axial center of the volume. In what follows, we refer to $I_{z,\Delta z_0}$also as “single-plane LFM images”.

Due to this acquisition geometry, direct summation over all depths z of $\{I_{z,\Delta z_0}\}$ would not form a synthetic LFM capture that contains sources across its entire depth range. To synthesize a volumetric LFM movie in which fluorescence appears to be emanating from across the entire depth range, we computationally shifted the apparent depth of the fluorescent source plane in the single-plane LFM movies and added up the results.

For example, to synthesize a volumetric LFM movie that covers the absolute depth range 100–300 µm (i.e., with the native focal plane at 200 µm), we started from the single-plane LFM movies recorded at absolute depths 100, 125, 150, …, 300 µm and shifted the fluorescent source planes contained in them such that they appeared at the correct axial offsets in the shifted light field data. In each frame of the single-plane LFM movie recorded at absolute depth 150 µm, the source plane appears at a relative offset of −40 µm from the native focal plane. In the output LFM dataset, this plane should appear at an offset of +50 µm relative to the native focal plane at 200 µm. Therefore, it needs to be shifted by +90 µm before being added into the output LFM movie.

The required light field shift operation can be derived by starting from the expression for refocusing the data contained in a light field capture to form the image that a non-light-field camera would capture were it focused to a specific depth in the volumetric FOV of the light field capture. This “refocusing” operation is e.g. formulated as follows in Ref. 22:

$$E_{(\alpha\cdot F)}(x',y') = \frac{1}{\alpha^2 \cdot F^2} \iint L_F^{(u,v)}(u(1-1/\alpha)+x'/\alpha\,, v(1-1/\alpha)+y'/\alpha)\,du\,dv$$

Here, $L_F^{(u,v)}(x,y)$ is the light field capture, represented as a set of sub-aperture images denoted by indices $(u,v)$ which enumerate the pixels behind each microlens in the canonical image-space LFM design[3] and correspond to individual perspective views on the source scene. $F$ denotes the distance between the exit pupil of the imaging lens and the image plane in the actual light field capture (and is fixed in our application). The refocusing factor $\alpha$ is the relative scaling in distance between the exit pupil and the virtual film plane at which to refocus the image. $E_{(\alpha\cdot F)}(x',y')$ is the resulting refocused image, with pixel coordinates in the virtual image plane denoted as $(x',y')$. The expression above corresponds to a dilation by a factor $\alpha$, followed by a shift by $\left(u\left(1-\frac{1}{\alpha}\right), v\left(1-\frac{1}{\alpha}\right)\right)$, and summation over all the sub-aperture images contained in the light field capture.

The above expression hints at a transformation of $L_F^{(u,v)}(x,y)$ that would lead to a given source depth F to be refocused at a different virtual film plane $(\alpha'\cdot F)$: It simply consists in an inverse shift of the sub-aperture images, with the factors $\alpha$ and $\alpha'$determining the axial shift distance.

We determined the factors $\alpha$ for each desired shift distance by calibration based on the numerically simulated PSF of our LFM: A 2D test image was convolved with the 3D LFM PSF, the resulting light field shifted with a series of factors $\alpha \in [-2,2]$, the resulting shifted light fields were each 3D-reconstructed by deconvolution with the LFM PSF, and the depth of best focus of the input image was determined in the resulting

LFM volume. Thus, a calibrated mapping between desired shift distance $z$ and corresponding shift factor $\alpha_z$ was established that is optimal for the PSF of our system.

To synthesize realistic volumetric data from our experimental LFM raw captures $I_{z,\Delta z_0}$ it was necessary shift $I_{z,\Delta z_0}$ to $I_{z,\Delta z'}$ where $\Delta z'$ could be different from $\Delta z_0$. If we denote the sub-aperture image with perspective indices $(u, v)$ of $I_{z,\Delta z'}$ as $I_{z,\Delta z_0}^{(u,v)}$, then the required transformation is

$$I_{z,\Delta z'}^{(u,v)} = I_{z,\Delta z_0}^{(u,v)}\left((-u + x)\left(\frac{1}{\alpha_{\Delta z_0}} - \frac{1}{\alpha_{\Delta z'}}\right), (-v + y)\left(\frac{1}{\alpha_{\Delta z_0}} - \frac{1}{\alpha_{\Delta z'}}\right)\right)$$

where $\alpha_{\Delta z_0}$ and $\alpha_{\Delta z'}$ are the values of $\alpha_z$ corresponding to the fixed offset $\Delta z_0$ to the native image plane and to at the variable shift $\Delta z' = z - z_{center}$. Here, $z_{center}$ is the depth of the native focal plane relative to the dura mater at which to synthesize the final LFM volume. In this work, we chose $z_{center}$ = 100, 200, or 300 µm. The set of sub-aperture images of the synthesized volumetric LFM movie is then $I_{syn}^{(u,v)} = \sum_{\Delta z'} I_{z,\Delta z'}^{(u,v)}$. This 4D quantity, a light field in sub-aperture representation, is then reshaped to the 2D raw light field capture representation by placing the sub-aperture pixels at the correct locations behind each microlens. We verified the correctness of the operation by shifting, adding and 3D-reconstructing LFM test images and found excellent agreement between expected and obtained shifted depths of sources.

These light field shift operations are carried out for each frame in the movies. The resulting synthesis of experimental single-plane 2pM–LFM recordings into an LFM volumetric dataset contains sources at all depths $z$ and serves as input to the MesoLF analysis pipeline. The outputs of MesoLF are then compared to the human-annotated outputs of CaImAn applied to the 2pM data to obtain MesoLF performance metrics (Methods, main text).

The light field shift was implemented as the MATLAB function `virtual_shift()` contained in Supplementary Software 1.
